# Respiratory syncytial virus load-dependent ciliated cell dedifferentiation and downregulation of immune response genes

**DOI:** 10.1101/2025.08.01.666372

**Authors:** Kevin Berg, Sibylle Haid, Ehsan Vafadarnejad, Arnaud Carpentier, Robert Geffers, Bettina Wiegmann, Emmanuel Saliba, Florian Erhard, Thomas Pietschmann

**Affiliations:** Institute of Virology and Immunology, Julius-Maximilians-University Würzburg, Würzburg, Germany; Faculty for Informatics and Data Science, University of Regensburg, Regensburg, Germany; Institute for Experimental Virology, Centre for Experimental and Clinical Infection Research, a Joint Venture between Helmholtz Centre for Infection Research and the Hannover Medical School, TWINCORE, Hannover, Germany; German Center for Lung Research (DZL); Helmholtz Institute for RNA-based Infection Research (HIRI), Helmholtz-Center for Infection Research (HZI), Würzburg, Germany; Genome Analytics, HZI, Braunschweig, Germany; Department of Cardiothoracic, Transplantation and Vascular Surgery, Hannover Medical School, Hannover, Germany; Lower Saxony Center for Biomedical Engineering, Implant Research and Development (NIFE), Hannover, Germany; Institute of Molecular Infection Biology Faculty of Medicine, University of Würzburg, Würzburg, Germany; German Center for Infection Research (DZIF), Site Hannover-Braunschweig, Germany

## Abstract

Respiratory syncytial virus (RSV) can cause severe lower respiratory tract infections in infants, older adults, and individuals with chronic or immunocompromising conditions. To understand how RSV affects airway cells, we infected primary human airway epithelial cultures with RSV and analyzed infected and bystander cells using time-resolved single-cell RNA sequencing and imaging. RSV mainly infected ciliated cells, triggering a virus load-dependent shutdown of genes involved in ciliogenesis, antigen presentation, and innate sensing, including key interferon (IFN) and pattern recognition pathways. Only a subset of infected cells produced type I and III IFNs, while bystander cells exhibited strong IFN-stimulated gene (ISG) signatures. Neither IFN treatment nor ISG induction eliminated infection, but IRF1, an antiviral transcription factor not suppressed by RSV, remained robustly expressed. Ectopic IRF1 expression *in vitro* reduced viral replication. These findings reveal how RSV evades antiviral defenses and highlight IRF1 as a potential target for therapeutic intervention.

## Introduction

The respiratory epithelium is a unique biological barrier with key functions in gas exchange and immune defense. Multiple cell types collaborate to fulfill these functions in a spatially organized fashion with multiciliated, goblet, secretory and basal cells as dominating cell types of the trachea and proximal airway ^1^. Upon pathogen exposure, the epithelium mounts an innate immune response, including the secretion of cytokines and chemokines that shape the local inflammatory milieu. These signals recruit and educate adaptive immune responses, ultimately clearing infections by humoral and cell-based effector mechanisms ^2^.

While these defense processes are ongoing, it is critical to maintain barrier functions and gas exchange. Therefore, immune silencing, differentiation, healing and regenerative processes are in place to maintain respiratory tract homeostasis during pathogen invasion, control and clearance. The recent COVID-19 pandemic has highlighted wide differences in the acute course of respiratory viral infections, the degree of lung damage and long-term effects. Furthermore, recent studies with single cell-resolved transcriptional profiling provide new insights into the mechanistic underpinnings of severe viral respiratory lung disease including involved cell types ^3^. Specifically, in SARS-CoV-2 infections, a dampened and ineffective IFN response in the respiratory epithelium was associated with severe courses of infection ^4^.

The respiratory syncytial virus (RSV) is a globally distributed virus causing episodes of epidemic spread in humans across all age groups mostly during winter seasons. In healthy adults, the course of infection is usually mild with a self-limiting involvement of the upper respiratory tract. However, preterm infants, children younger than six months of age, and adults older than 60 years, as well as people with comorbidities such as for instance chronic lung or heart diseases as well as immune-deficient or immune-suppressed are at risk for severe courses of acute lower respiratory tract infections.

The conditions for preventing severe courses of RSV infection have much improved in 2023 due to the approval of prophylactic vaccines and the licensing of the monoclonal antibody nirsevimab ^5^. However, both currently available prophylactic antibodies have an efficacy between 40% to 80% ^6,7^ and the vaccines for older adults an efficacy between 66.7 and 82.6% ^5^. Moreover, treatment options for RSV infection are limited to symptomatic therapy. Therefore, novel therapeutic approaches and antiviral targets including host-targeting strategies should be explored.

The mechanisms of RSV-associated lung disease are diverse, but likely include direct effects of RSV on infected lung cells. Immunohistochemical analyses of lung tissue from fatally infected infants and analyses of human cell cultures of primary lung epithelial cells show that RSV primarily, but not exclusively, infects ciliated epithelial cells ^8–13^. Tissue sections also revealed infected type 1 and 2 pneumocytes, and occasionally other non-ciliated infected cells ^9,13^. Histochemical staining of lung tissue from RSV-infected neonatal baboons and newborn lambs has provided evidence that basal cells are also infected by RSV *in vivo*, possibly following airway injury due to RSV-induced shedding of infected ciliated cells ^14,15^. Consistent with this, *in vitro* data in primary human bronchial epithelial cells (HBEC) cultured under air-liquid interface (ALI) conditions show that sub-apical basal cells are also infected, at least after injury of the pseudostratified lung epithelium. In this model, it has been shown that infection of the basal cells leads to the production of interferons, which in turn influence the differentiation of the cells to produce more secretory and fewer ciliated cells ^16^. The authors of this study hypothesized that this could be a mechanism to explain reduced numbers of ciliated cells reported in RSV-infected humans ^16,17^. In addition to affecting basal cell differentiation, apoptosis and sloughing of RSV-infected ciliated cells, and occasionally observed syncytia formation, as documented in well-differentiated primary paediatric bronchial epithelial cells (WD-PBECs), are likely to explain this phenomenon and the pathogenesis of RSV *in vivo* ^12,18^.

At least 3 out of the 11 viral proteins, namely the non-structural proteins NS1 and NS2 and the glycoprotein G, interfere with the type I interferon (IFN) response ^19–23^. Nevertheless, RSV infection elicits an inflammatory response by the activation of retinoic acid-inducible gene I (RIG-I) and melanoma differentiation-associated protein 5 (MDA5) and the Toll-like receptors TLR-2, TLR-3, TLR-4 and TLR-7 ^24,25^, leading to the secretion of cytokines like IL-6 and IL-8 and type I and III interferons. Recent studies emphasize that the host’s innate immune state - both before and during RSV infection - plays a critical role in determining infection outcome. Transcriptional profiling in blood, nasal tissue, and airway epithelial cells has revealed that variations in immune activation can influence susceptibility, viral replication, and disease severity ^26–29^. These findings underscore the importance of understanding early immune responses to RSV at the tissue and cellular level.

Therefore, here we analyzed the kinetics and characteristics of host cell responses to RSV in an adult primary human pseudo-stratified airway epithelial cell model. We chose this model because it is based on primary human cells, includes multiple lung cell types, and recapitulates important features of RSV-host interactions in the human lung ^12^. We performed scRNA-seq on air-liquid interface cultures from six donors at 1, 3, 5 and 7 days post infection. Our analysis focused on transcriptional dynamics across the viral load gradient, revealing a continuous host-transcriptional change that characterizes the host response along the progressing RSV replication cycle at the single-cell level. As viral load increased, ciliated cells progressively lost their defining markers and transitioned to a basal-like cell state. In addition, increasing viral load was associated with a waning type I IFN innate immune response, decreased expression of MHC class II subunit transcripts, and an increased cellular stress response, potentially leading to apoptotic cell death. However, we also found that selected interferon-stimulated genes (ISGs) such as interferon regulatory factor 1 (IRF1), a potent transcription factor that controls numerous antiviral host genes, were not downregulated by RSV infection. Furthermore, we demonstrated that IRF1 overexpression, unlike exogenous addition of type I IFN, significantly reduced RSV infection, suggesting its potential as a prophylactic and therapeutic intervention for RSV infection.

## Results

### RSV infection elicits distinguishable transcriptional responses in infected and bystander cell populations

To study RSV infection and host responses by single-cell RNA-seq in a model system that mimics the human *in vivo* situation, we cultured primary human airway epithelial cells in air-liquid-interface (ALI) cultures from six donors and infected them with a GFP-expressing RSV-A strain Long reporter virus with 1.4×10^5^ TCID_50_ per well (Methods, **Figure 1A**). ALI cells were dissociated and sorted by fluorescence activated cell sorting (FACS) for GFP expression prior to scRNA-seq to distinguish between RSV infected (GFP-positive, ranging from 0.7 to 3.2%, **Supplementary Figure S1A-B**) and bystander cells (GFP negative) on days 1, 3, 5 and 7 post infection. In parallel, we also analyzed mock cells (from the same six donors) that were not exposed to RSV but treated with conditioned media as control. This strategy allows to enrich the otherwise only weakly represented infected cells. Pools of GFP-positive, GFP-negative and mock cells were loaded on the 10x Genomics controller and *in silico* demultiplexed using hashtags for time points and genetic background to differentiate the donors (**Figure 1A**). Altogether 12,331 cells (mock, n=6,496; Infected, GFP-positive, n=2,037; Bystander, GFP negative, n=3,798) with an average of ∼5,700 genes per cells met quality control criteria and were further analyzed (see Methods, **Supplementary Figure S2A-D**).

**Figure 1:**
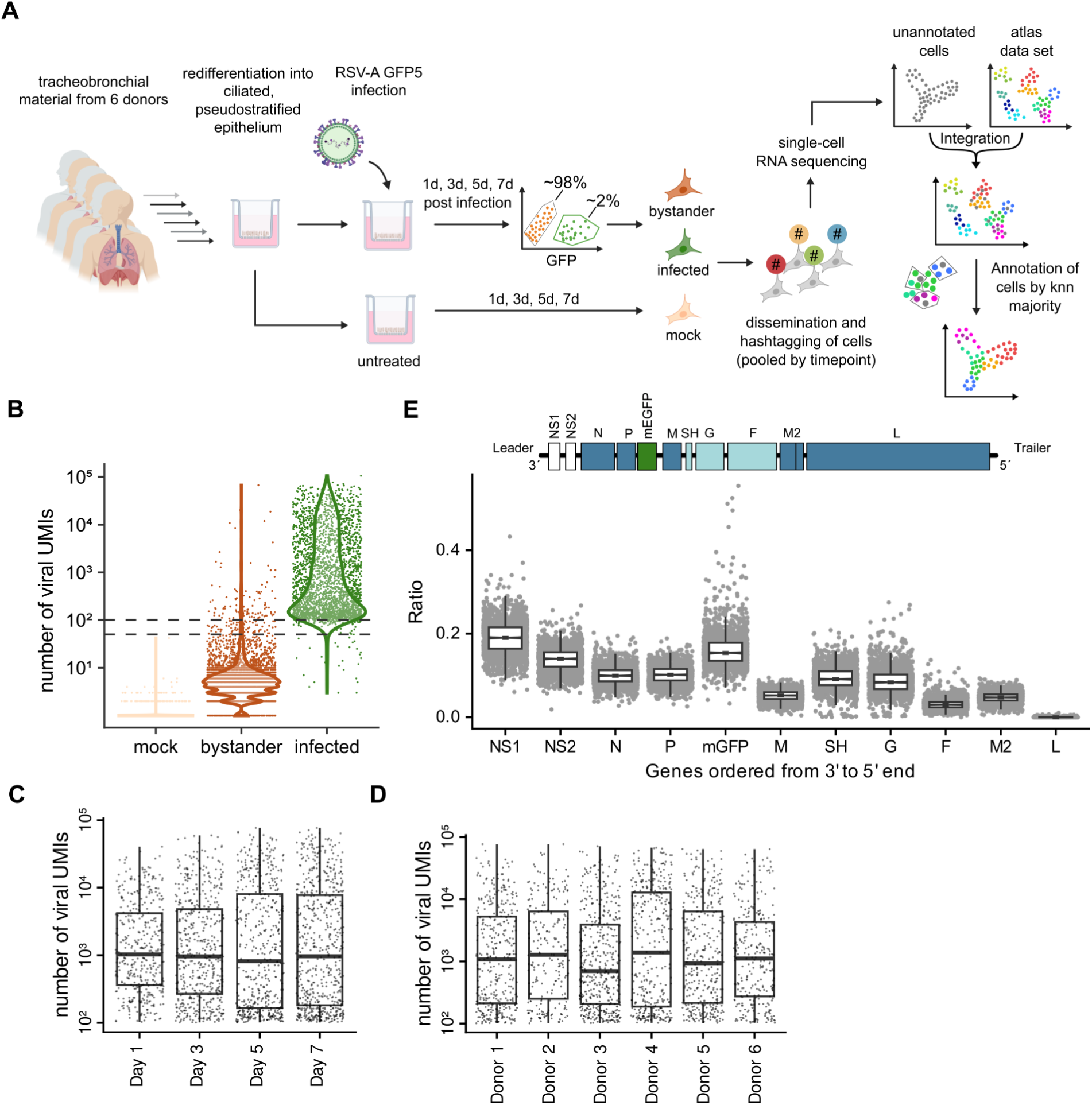
Study design and viral gene expression of RSV infected cells. (A) Tracheobronchial samples from 6 donors were cultured in ALI and infected with a GFP expressing RSV A strain. The cells were collected and fluorescence-sorted at days 1, 3, 5 and 7 post infection, subsequently hashtagged with antibodies and analyzed by scRNA-seq. Cell type annotation was done by integration of the data set with an atlas data set, calculation of the nearest neighborhood graph and annotation by majority of cell types in a cell’s neighborhood (see Methods, and Figures 2 and 3). (B) Violin plot showing the number of viral UMIs per cell. The dashed lines indicate the upper filter for bystander cells at 50 UMIs and the lower filter for infected cells at 100 UMIs. (C) Box plots of the viral UMIs at 1, 3, 5 and 7 days post infection. (D) Box plots of the viral UMIs across 6 donors. (E) Transcriptional profile of viral mRNA after manual RSV reannotation. The viral genome is given at the top and the position of the GFP reporter gene is annotated.

First, we focused on distinguishing the infected from bystander cells. Viral mRNA copy numbers differed by more than 1,000-fold among infected cells with detectable GFP signal, highlighting the massive heterogeneity in infection (**Figure 1B**). Consistent with FACS sorting, viral RNA copy numbers were well separated between infected and bystander cells. Most bystander cells had viral RNA loads of less than 10 detected mRNAs (counts of unique molecular identifiers, UMIs), an order of magnitude lower than the majority of infected cells. Very few bystander cells had viral RNA copies greater than 50 UMIs (n=173, 2.89% of all bystanders), which may be caused by RSV particles adhering to these cells and/or RSV RNA released from infected cells binding to these cells, or by FACS errors. In order to clearly distinguish the transcriptional profiles of bystander and infected cells from each other, we only included bystanders below 50 viral UMIs and infected cells above 100 viral UMIs in our subsequent analyses. Interestingly, the distribution of viral UMIs among GFP-positive cells was very similar across time points as well as donors with a median of roughly 1,000 UMIs and UMI counts spanning almost three orders of magnitude, which is equivalent to <0.5% and ca. 56% of total cellular RNA, respectively (**Figure 1C-D**). From the perspective of a single GFP-positive cell, viral gene expression is expected to increase over the course of time and the viral infection cycle. Thus, the observed similarities in viral RNA load between the individual time points suggest that also at day 3 to 7, there are cells that have experienced a recent onset of viral gene expression, and others that have already reached the maximum viral RNA load that the cells can sustain. This observation suggests that the virus replication cycle in this model is completed in less than 24 hours and that there are continuous rounds of infection and reinfection over the course of 7 days. Notably, this was observed consistently for all six donors (**Supplementary Figure S3**).

Consistent with previous reports ^30–32^, viral mRNA expression decreased from the 3’ to the 5’ end of the viral genome, with the most 3’ transcription unit encoding the NS1 protein being the most abundant and the most 5’-terminal message of the L protein being the least transcribed mRNA (**Figure 1E**). Interestingly, in our initial analysis of this gradient, the G protein mRNA was clearly over-represented relative to its position within the transcriptional gradient (**Supplementary Figure 4A**). On closer inspection of the underlying read mapping pattern to the RSV reference genome ^33^, we found two irregularities. First, for four genes (mGFP, M, SH, F), reads were not fully covered by the reference annotation, owing to a lacking annotation of 3’ UTRs in the reference (**Supplementary Figure S4B**). Second, and unique to the G protein sequence, we observed a second peak of mapped reads in the center of the gene body (**Supplementary Figure S4B**), which substantially increased the overall G mRNA read counts. Examining the G gene sequence, we found several polyA stretches downstream of the second peak within the G protein coding sequence, similar in distance to the polyA signal and the primary peak (**Supplementary Figure S4C**). During reverse transcription, these additional polyA sites could provide additional anchor points for polyT primer binding, resulting in an overrepresentation of the G protein mRNAs. Taken together, both the shortened 3’ UTRs and the internal peak in the G protein message led to biased read counts for viral mRNAs. To address both irregularities, we instead created a manual, data-driven annotation of the RSV genome for the analysis of the transcriptional gradient, using fixed 200bp read count analysis windows centered on the 3’ peaks for each gene. Using this curated annotation (**Supplementary Figure S4D**), still an imperfect 5’ to 3’-polar mRNA gradient is apparent. The overrepresentation of the mGFP, SH, G and M2 messages relative to their respective genome position (**Figure 1E**) suggests distinct post-transcriptional regulation of these mRNAs.

### Mock gene expression profiles reflect the major epithelial cell types but infection globally perturbs gene expression

To investigate RSV infection at the single cell level, we transferred cell type annotations from a published human lung cell atlas ^34^ first to our mock cells (**Figure 2A**). Label transfer was favored over marker gene based manual annotation of cell types to make use of the comprehensive lung cell atlas that has been annotated by large gene panels (see Methods). Ionocytes and Club cells were filtered out because only a handful of them could be captured (< 10 cells). We identified a distinct cluster of 198 cells enriched with low quality that was also filtered out (**Supplementary Fig. 2B-C**). Furthermore, we were able to assign basal and ciliated cell subpopulations (e.g. proliferating basal cells) identical to those previously assigned by the human lung cell atlas. This cell type assignment was overall conserved across all six donors (**Supplementary Fig. 2F**). Thus, our culture model faithfully recapitulated the typical cell types of a pseudo-stratified human bronchial epithelial cell layer ^34^.

**Figure 2:**
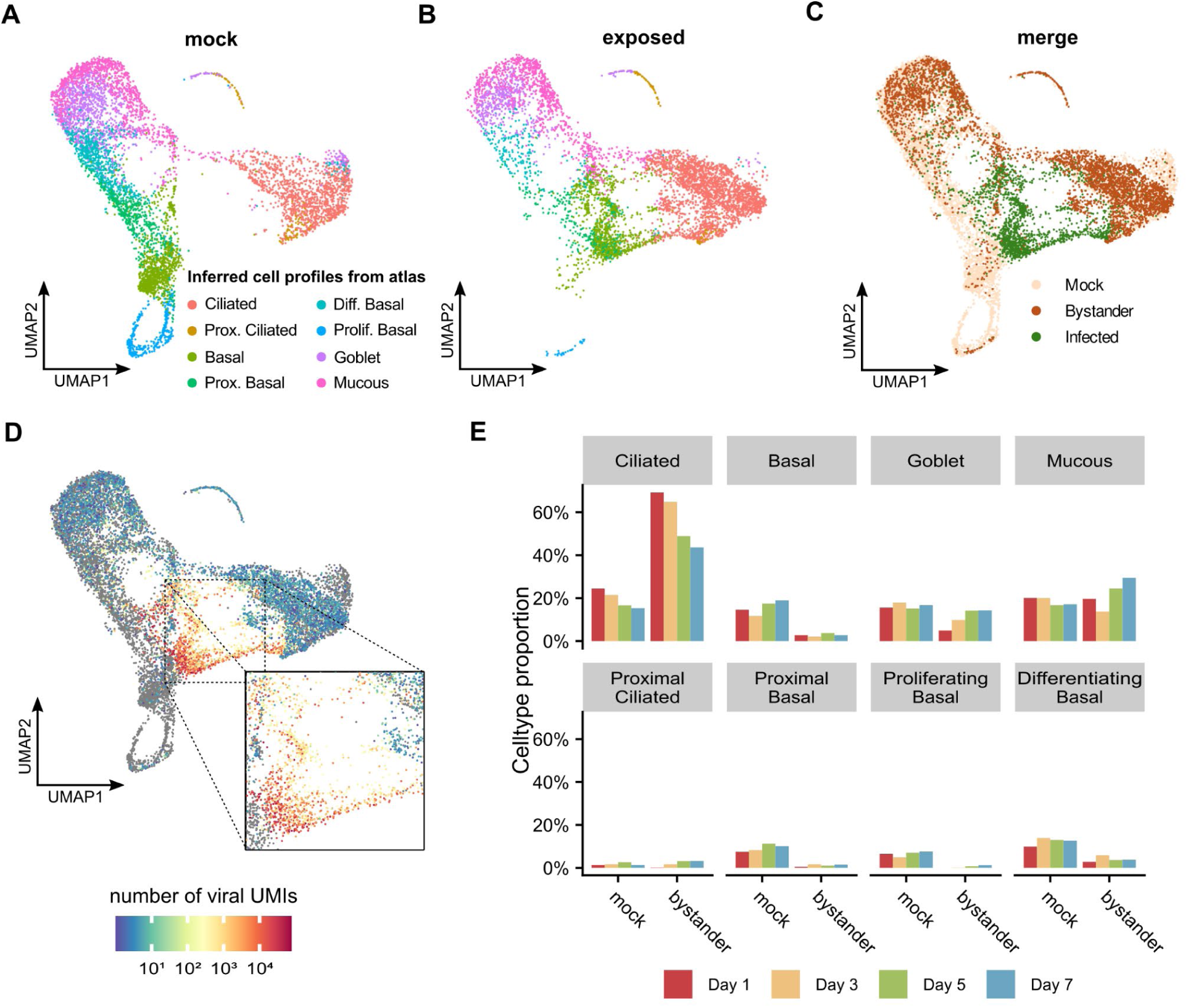
Cell profile annotation via human lung cell atlas. (A) UMAP representation of mock cell profiles annotated by integration with human lung cell atlas. (B) UMAP representation of RSV exposed cell profiles annotated by integration with human lung cell atlas. (C) UMAP representation of infection states of mock and exposed cells. (D) UMAP representation of the number of viral UMIs across infected cells. The inlay displays a magnified view of highly infected cells. (E) Bar plots of the relative proportion of cell types within mock and bystander cells across time points. The x-axis represents the different infection states, the y-axis indicates the relative proportion of the cell type per time point and infection state, with the total proportions of all cell types per infection state and at each time point summing to 1.

Next, using canonical correlation analysis, we mapped RSV exposed cells including bystanders and infected cells into a joint embedding with the mock cells for an automatic cell type annotation of the exposed cells (**Figure 2B-C**). Overall, exposed cells matched well to mock cells with two exceptions (**Figure 2A-C**): First, all basal cell types (basal and proximal, proliferating and differentiating basal) were strongly underrepresented among exposed and absent from infected cells, presumably because the differentiation of basal cells is strongly induced upon depletion of cells by infection; second, the majority of infected cells segregated away from all mock cell subsets and shared expression patterns of both ciliated and basal mock cells in a viral RNA load dependent manner. Infected cells with low viral RNA load appeared most similar to ciliated cells and were thus called “ciliated-like” (**Figure 2D**). By contrast the infected cells with highest viral RNA load were rather correlated with basal cells and therefore called “basal-like”. Importantly, this phenomenon was consistently observed throughout the 4 days investigated here and for all six donors (**Supplementary Fig 3**).

To further investigate these phenomena, we examined transferred cell type annotations and compared quantitative changes of cell populations over time in mock and bystander cells. In mock-treated control cells, the relative numbers of different cell types remained fairly constant over day 1 to 7 with only a slight decrease in ciliated cells and a slight increase of basal cells over time, indicating that the cell culture is in a steady-state equilibrium when unperturbed (**Figure 2E**). In contrast, the distribution of cell types was significantly altered in the RSV-exposed bystander cell populations, with an almost complete loss of proliferating basal cells, a significantly decreased proportion of proximal and differentiating basal cells, a strong overrepresentation of ciliated (or ciliated-like cells), and a time-dependent modulation of the ciliated, goblet, and mucous cell compartments. Ciliated cells initially increased and returned towards baseline levels by day 7, whereas goblet cells initially decreased and returned to levels similar to mock-treated controls by day 7 and mucous cells increased at day 5 and 7 (**Figure 2E**). Taken together, these results showed that RSV infection caused an expansion of the ciliated cell compartment and did not deplete ciliated cells or completely spread to all available ciliated cells. In contrast, exposure to RSV severely depleted basal cells relative to their abundance in mock-treated controls.

### RSV primarily infects ciliated cells and causes cell de-differentiation

Previous studies in similar culture models have reported that RSV primarily targets ciliated cells ^10–12,16,35^. Likewise histological analyses of RSV-infected pediatric lungs point towards ciliated cells as primary RSV target cells ^9,13^. It was therefore surprising that the majority of RSV-infected cells with high viral RNA load had gene expression profiles reminiscent of basal cells and were clearly distinct from ciliated cells (**Figure 2A-D**).

To investigate the alterations in the gene expression profiles of infected cells, we first dissected cell type specific marker gene expression in the mock, bystander and infected cell populations (**Figure 3A**). Mock cells comprised well-distinguishable cellular mRNA expression signatures separating ciliated and basal cells from each other. Very high expression of *FOXJ1*, *ARL13B*, *CFAP53* and *RSPH9* characterized essentially all ciliated cells. In contrast, the vast majority of basal cells abundantly expressed *ITGA6*, *TP63*, *NGFR, KRT5*, *KRT13*, *KRT15* and *KRT17* mRNAs (**Figure 3A**, top panel). Interestingly, with the exception of *KRT5* and *KRT13*, all markers showed virtually the same expression patterns in the respective bystander cells.

**Figure 3:**
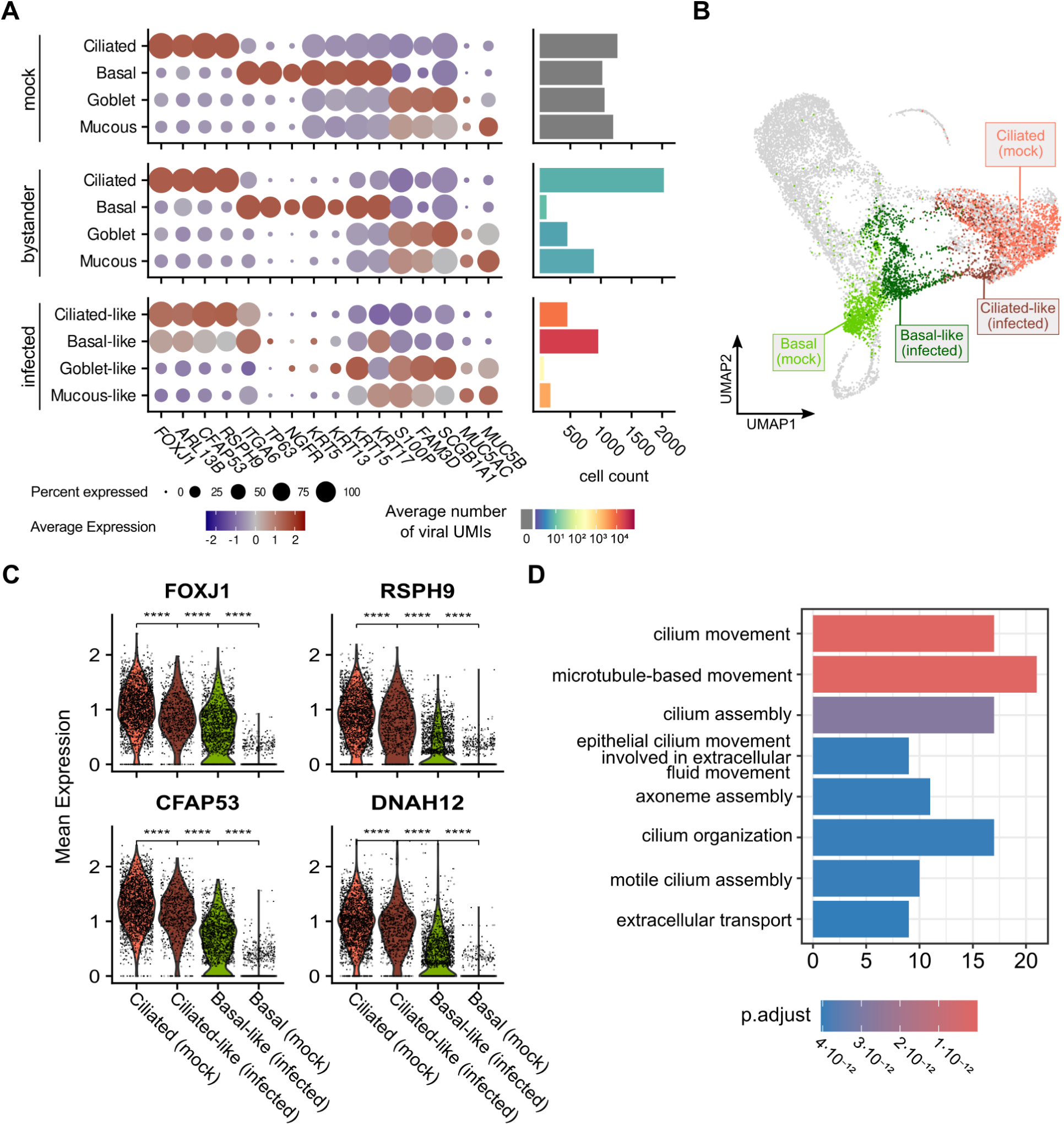
Infected ciliated cells with high virus load exhibit basal cell-like expression profile. (A) Average expression and percentage of expressing population of marker genes in mock, bystander and infected ciliated, basal, goblet and mucous cells (left). Cell counts per cell type and infection state, the color represents the average natural log value of viral reads (right). (B) UMAP visualization of basal(-like) and ciliated(-like) cells. (C) Gene expression of 4 marker genes for ciliated cells and microtubules over ciliated, ciliated-like, basal-like and basal cells. Wilcoxon test, p-values indicated. (D) Gene ontology analysis of downregulated genes in infected Basal-like cells versus mock ciliated cells. The bars represent the gene count per category, the color represents the respective adjusted p-value. The top 8 hits are shown.

By contrast, infected cells showed a strongly perturbed pattern of marker gene expression: In the ciliated-like cells, which were characterized by low viral RNA load, the expression of *FOXJ1*, *ARL13B*, *CFAP53* and *RSPH9* was slightly reduced compared to mock ciliated cells. These ciliated cell marker genes were expressed even more weakly in the highly infected basal-like cells. This indicated that RSV reduced the expression of these genes over the course of its infection cycle. In contrast, the basal cell marker gene *ITGA6* was more strongly expressed in ciliated-like infected cells than in mock ciliated cells, and showed even stronger expression in basal-like infected cells.

We also detected a few goblet-like and mucous-like infected cells with very low or low virus load, respectively. Again, their transcriptional profile was skewed compared to the respective cell populations in the mock-treated culture. These latter changes are most prominently represented by the upregulation of *KRT15* in the goblet-like infected cells compared to the goblet cells and the upregulation of *KRT17* in the mucous-like cells compared to the mucous cells. Collectively, these results indicate that RSV infection induces a profound transcriptional reprogramming that affects cell annotation and likely cell physiology. Using viral RNA load as a marker of the infection time of an individual cell, we hypothesized that ciliated cells were infected throughout the seven days studied here and changed their gene expression profile towards a more basal cell-like profile as its infection cycle progressed, suggesting that rising virus RNA load may drive dedifferentiation of ciliated cells towards a basal-like phenotype.

To investigate this dedifferentiation in more detail, we compared the infected and mock cells annotated as “ciliated” or “basal” at the single cell level (**Figure 3B**). All four ciliated marker genes *FOXJ1*, *ARL13B*, *CFAP53* and *RSPH9* were most strongly expressed in ciliated mock cells (**Figure 3C**). In infected ciliated-like cells with low viral RNA load, ciliated marker gene expression was markedly reduced. This reduction was even stronger in infected basal-like cells with higher viral RNA load. Gene ontology analysis of the cumulated transcriptional changes distinguishing the infected basal-like cells from the ciliated mock cells identified a strong and significant under-representation of GOs involved in cilium assembly, organization and movement but no other pathway that is unrelated to cilia (**Figure 3D**). These observations are consistent with a model where ciliated cells are infected by RSV, which then lose their ciliated phenotype during the infection cycle.

To confirm ciliated cells as the primary target cells of RSV in this culture system and to investigate the fate of these cells, we analyzed RSV infection by confocal microscopy and flow cytometry using antibodies against proteins that distinguish ciliated from basal cells (**Figure 4**). Generally, basal cells were localized to the basolateral layer whereas ciliated cells localized to the apical side. Interestingly, most infected cells were localized to the apical surface of the culture. However, only a few infected cells were clearly decorated with prominent tubulin staining at their apical surface, indicating cilia and infection of ciliated cells by RSV. Moreover, very few infected cells co-expressed Cytokeratin 5/KRT5, which is characteristic of basal cells. Our flow cytometry analysis supported the observations of scRNA profiling and documented that appr. 95% of infected cells co-express ARL13, a marker for ciliated cells, whereas only about 10 percent express KRT5. Taken together, these observations indicate that RSV primarily initiates viral gene expression in ciliated cells and causes a profound, and virus-load dependent de-differentiation of these cells.

**Figure 4.**
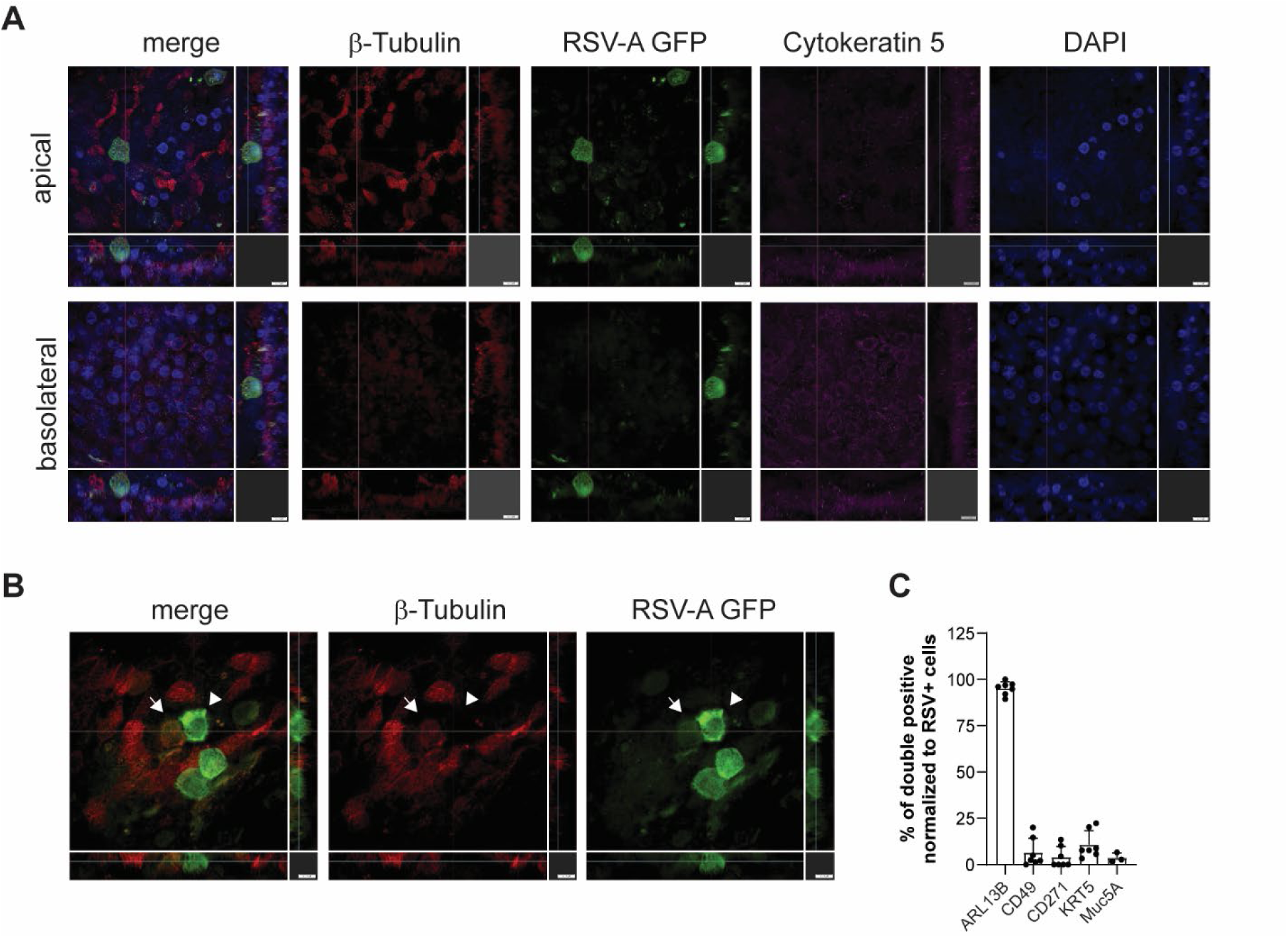
RSV primarily infects ciliated cells. (A) Immunofluorescence analysis of well-differentiated primary human airway epithelia cells cultured under ALI conditions and infected with an RSV-A GFP reporter virus. ß-Tubulin primarily stains the cilia whereas Cytokeratin 5/KRT5 is a marker for basal cells. Two different projections (upper row: apical, lower row: basolateral) as well as z-stacks are displayed. Scale bar: 100 µm. One representative set of pictures is given. (B) Immunofluorescence analysis of primary human airway epithelial cells infected with RSV-A-GFP reporter virus. ß-Tubulin staining is reduced in cells with high GFP expression (compare b-Tubulin staining in cells with weak GFP expression (arrow) with cells with strong GFP expression (arrow head). (C) Flow cytometric analysis of well-differentiated human airway epithelia cells. Percentage of RSV-infected cells positive for respective marker protein is depicted. Symbols represent individual donors from n=2 independent experiments.

### RSV mounts a strong IFN response that is quickly blunted during infection

We observed that the overall expression patterns of infected cells were very similar across the 4 experimental time points (**Figure 1C**). This allowed us to pool infected cells across all time points to further define RSV-induced mechanisms of cellular reprogramming. We ranked the infected cells according to their viral RNA load and divided these cells into 20 different pseudo-bulks, each containing the same number of cells (**Figure 5A**). These pseudo-bulks of infected cells thus represented a continuum from the lowest viral RNA load in class 1 to the highest represented by the cells in class 20. Principal component analysis of the 20 pseudo-bulk groups revealed gradually changing and well-differentiated transcriptional profiles that differed from both mock-treated ciliated and bystander cell profiles (**Figure 5B**). Since viral RNA levels are expected to rise upon infection of a single cell, we considered the virus RNA load a proxy for “infection time” representing how far a cell has progressed in the infection cycle (**Figure 5CD, Supplementary Fig. S3**).

**Figure 5:**
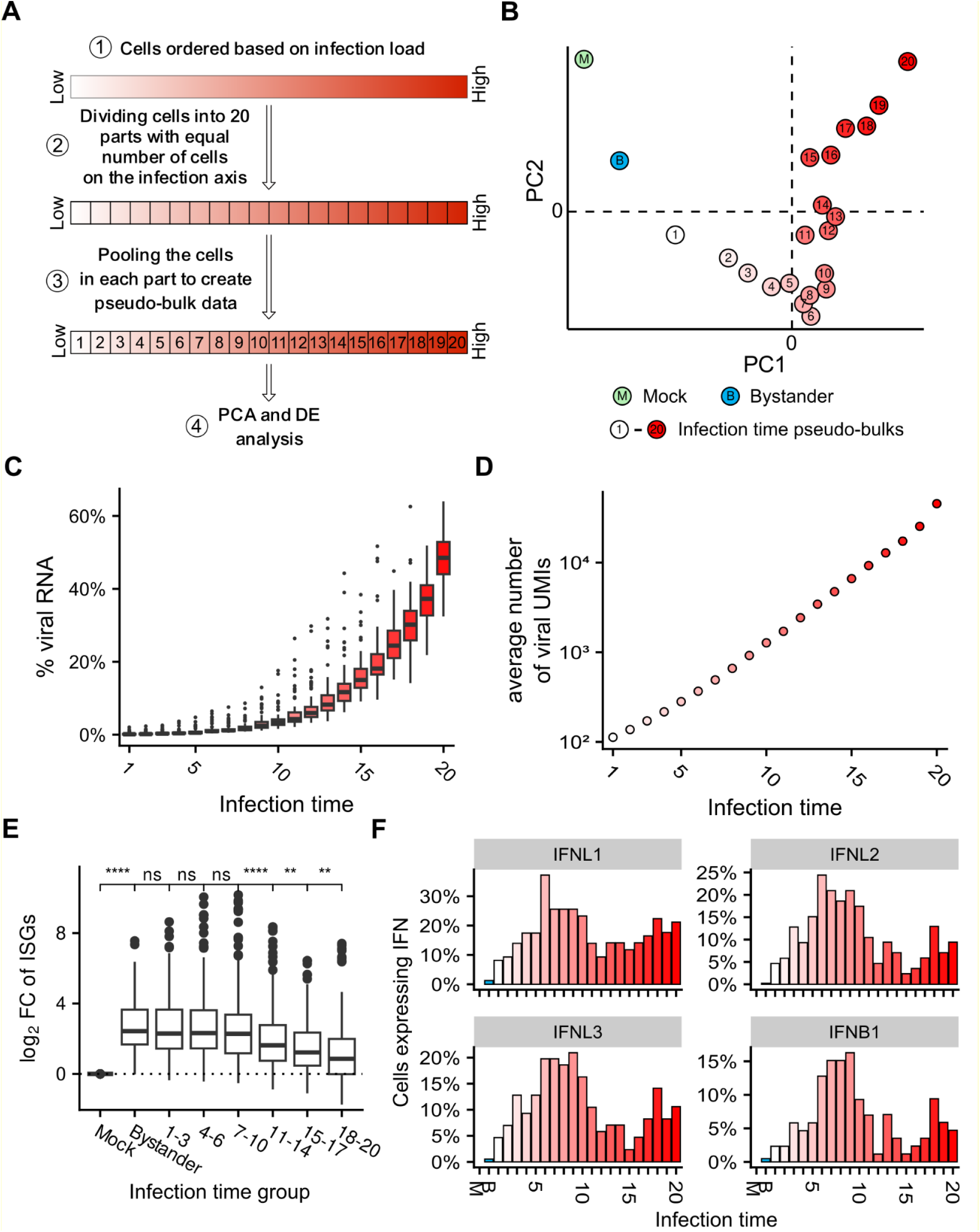
Pseudobulk analysis reveals infection time-dependent blunting of the interferon response. (A) Scheme of the pseudo bulk analysis to investigate viral load dependent transcriptional regulation. Cells are equally divided along the viral load axis into 20 bins, which are then subjected to PCA and differential gene expression analyses. (B) PCA plot of the first two principal components of the binned reads for mock (green), bystander (blue) and 20 viral load bins (red gradient). (C) Boxplots showing the percentage of viral RNA for all infected cells per infection time pseudobulk. (D) Average number of viral UMIs per infection time pseudobulk. (E) Visualization of the interferon score (mean LogFC of interferon hallmark genes) across mock and bystander cells and grouped viral load bin. Wilcoxon test, p-values indicated. (F) Percentage of cells expressing IFNL1-3 or IFNB1 over mock, bystander and infection time pseudo bulks.

RSV infection induces an interferon response in the human lung ^36–40^, in primary human cells ^41,42^, and in animal models ^43^, although IFN responses in RSV infection of human cells may be weaker than in influenza A virus infection ^36,41^. Such a comparatively weak interferon response to RSV may be due to the RSV proteins NS1 and NS2 blunting IFN induction and signalling ^44,45^. To quantify RSV-induced ISG induction, we thus examined a predefined set of typical ISGs and computed an ISG score reflecting the overall activation of the interferon pathway per cell. Comparing this ISG score among ciliated mock cell populations, all bystander cells and across the 20 infection time pseudobulks revealed a strong interferon response in bystander cells and a stable ISG expression throughout infection time 1 to 10 (**Figure 5E**). Thereafter, the strength of the interferon response decreased rapidly and continuously at later infection time points. Thus, with increasing viral RNA load, interferon signaling was attenuated, probably due to NS1- and NS2-dependent interference with interferon signaling in the infected cells ^44,45^. Analysing the IFN response over the seven days of infection in bystander cells revealed a burst of ISG induction on day 1 and subsequent maintenance of a continuously high IFN score until day 7. Moreover, virus load classes above 10 exhibited a decreased ISG induction compared with lower virus load classes across all time points (**Supplementary Fig. S5**).

We detected mRNA expression of a type I IFN (IFN-β) and three type III IFN (IFN-λ1, IFN-λ2, IFN-λ3) in 10-20% of the cells from each infection time pseudobulk (**Figure 5F**). The temporal expression pattern was highly similar among these four IFN mRNAs: They were not detected in mock cells, barely detectable in bystander cells and increased their expression from infection time 1 to 6. At infection time 10 and higher the number of type I and III IFN expressing cells was clearly reduced. Consistent with these data, we observed viral RNA load-dependent regulation of the pattern recognition receptor (PRR) mRNAs IFIH1 (MDA5), DDX58 (RIG-I) and TLR-3, but not IRF3 (**Figure 6A**). While the mRNAs of IFIH1, DDX58 and TLR3 were strongly induced in bystander cells and early infection load classes up to class 8, their expression decreased thereafter. Furthermore, with the exception of pseudobulk 15, where IFN expression was generally weakest (**Figure 5F**), expression of both IFIH1 and DDX58 correlated well (R>0.33, p<0.05, t test, Bonferroni corrected) with IFN expression, whereas a correlation of TLR3 was only present in pseudobulks of cells with low viral RNA load (**Figure 6B**). In contrast, the mRNA of IRF3 slightly increased in infected cells and was maintained at comparable levels across all viral RNA load classes. These results are consistent with NS1 and NS2 inhibiting the induction of type I and III IFNs and ISGs ^44,45^, including the mRNAs of key PRRs, which likely limits the ability of infected cells to induce IFNs. Notably, even at very low viral loads, only a fraction of RSV-infected cells had detectable levels of type I and III IFNs (approximately 5-10% of pseudobulk class I cells), suggesting that RSV infection is initially unrecognised in the majority of infected cells. To explore potential factors responsible for this, we quantified the mRNA expression heterogeneity of key PRRs in mock-treated ciliated cells (**Figure 6CD**). Notably, less than 25% of these cells expressed detectable DDX58 mRNA. Furthermore, DDX58 mRNA levels were among the most variable 5% of all detected mRNAs, suggesting that differential availability of PRRs in the early phase of infection may determine whether viral patterns can be sensed and IFNs are induced.

**Figure 6:**
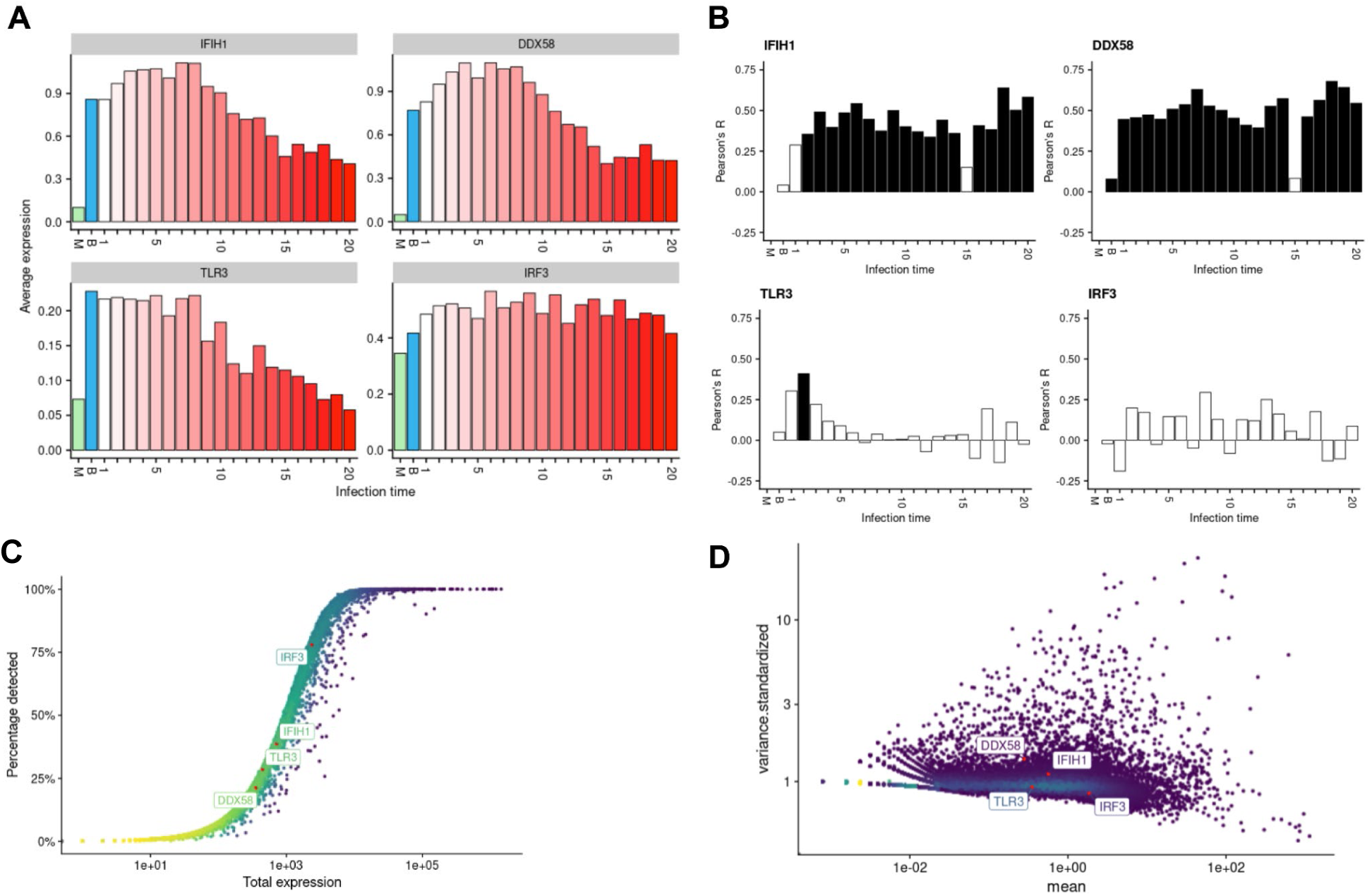
Regulation of PRR mRNAs in mock ciliated, bystander and RSV infected cells. (A) Average mRNA expression of given PRRs and IRF3 in mock, bystander and infected cells. Expression in infected cells was quantified across the gradient of virus load. (B) Correlation analysis between mRNA expression of IFIH1 (MDA5), DDX58 (RIG-I), TLR3 (TLR-3) and IRF3 type I and III IFN expression among cells for each pseudobulk. Black bars indicate statistical significance (p<0.05, t test, Bonferroni correction). (C) Relative number of mock treated cells (in %) expressing a given mRNA is displayed on the y-axis and total average expression of the given mRNA on the X axis. (D) The variance of in total 24,800 mRNAs detected in the pool of mock-treated ciliated cells, standardized by the respective average expression value (standardized variance) is plotted relative to the mean total expression. The DDX58 mRNA ranks at 1,034 and thus within the top 5% most variable mRNAs.

Since RSV substantially dampens the IFN response we wondered whether the fate of infected cells and the outcome of infection could be influenced by exogenous addition of type I IFN (**Supplementary Fig. S6**). However, exogenous supplementation of IFNα did not reduce the number of RSV infected cells, likely because RSV has established sufficient interference with IFN signaling to accommodate its infection and prevent clearance. Notably, addition of ruxolitinib, an inhibitor of STAT1-signaling, strongly increased the number of RSV infected cells (**Supplementary Fig. S6**).

In conclusion, ordering cells according to viral RNA load as a proxy for infection time of individual cells highlights the induction of IFN and its quick and highly efficient counterregulation by RSV at later infection times. Moreover, improved control of RSV would likely require enforcement of cellular antiviral defenses mechanisms that are not effectively blunted by RSV.

### RSV infection causes virus load-dependent transcriptional re-programming

Since both cell dedifferentiation **(Figure 3**) and interferon response (**Figure 5,6**) showed a viral RNA load-dependent progression of gene regulation, we set out to comprehensively investigate gene expression changes over the infection time defined by increasing viral RNA load. To this end, we compared gene expression in the 20 pseudobulk groups of infected cells with the one in the ciliated cells of the mock-treated control. In total, we detected 1,033 differentially expressed genes among all pseudobulk groups (1-20) of RSV infected cells and also in the mock-treated ciliated control cells. Clustering genes into eight groups according to their mRNA expression profile over the pseudobulk groups (**Figure 7A** and **Supplementary Fig. S7**) revealed distinct gene regulatory patterns. There were clusters showing a continuous down-(cluster 0) or progressive down-regulation (cluster 1) relative to mock-treated ciliated cells. Other clusters showed consistent (cluster 4) or progressive (cluster 6) gene up-regulation over infection time, or were characterized by a strong (cluster 2) or weak (cluster 3) initial gene expression burst coincident with the onset of viral gene expression and subsequent dampening. Finally, cluster 5 included genes whose expression was only weakly regulated across infection time, and cluster 7 included genes that were unchanged at low and high viral loads, but were slightly downregulated in the intermediate viral load pseudobulks. To further examine these distinct profiles and investigate whether they indeed represent separated clusters rather than a continuum of patterns, we developed a novel approach to visualize the progression of all differentially expressed genes over the infection cycle (defined by increasing viral RNA load). Typically, UMAP is used to embed cells in gene expression space, i.e., cells are placed in close proximity if and only if their gene expression profiles are similar. Here, we use UMAP to embed genes in a temporal space across the infection time. In other words, genes are placed in close proximity in this gene UMAP if and only if their expression pattern was similar over the course of the infection cycle, i.e. over the 20 pseudobulk groups (**Figure 7A**).

**Figure 7:**
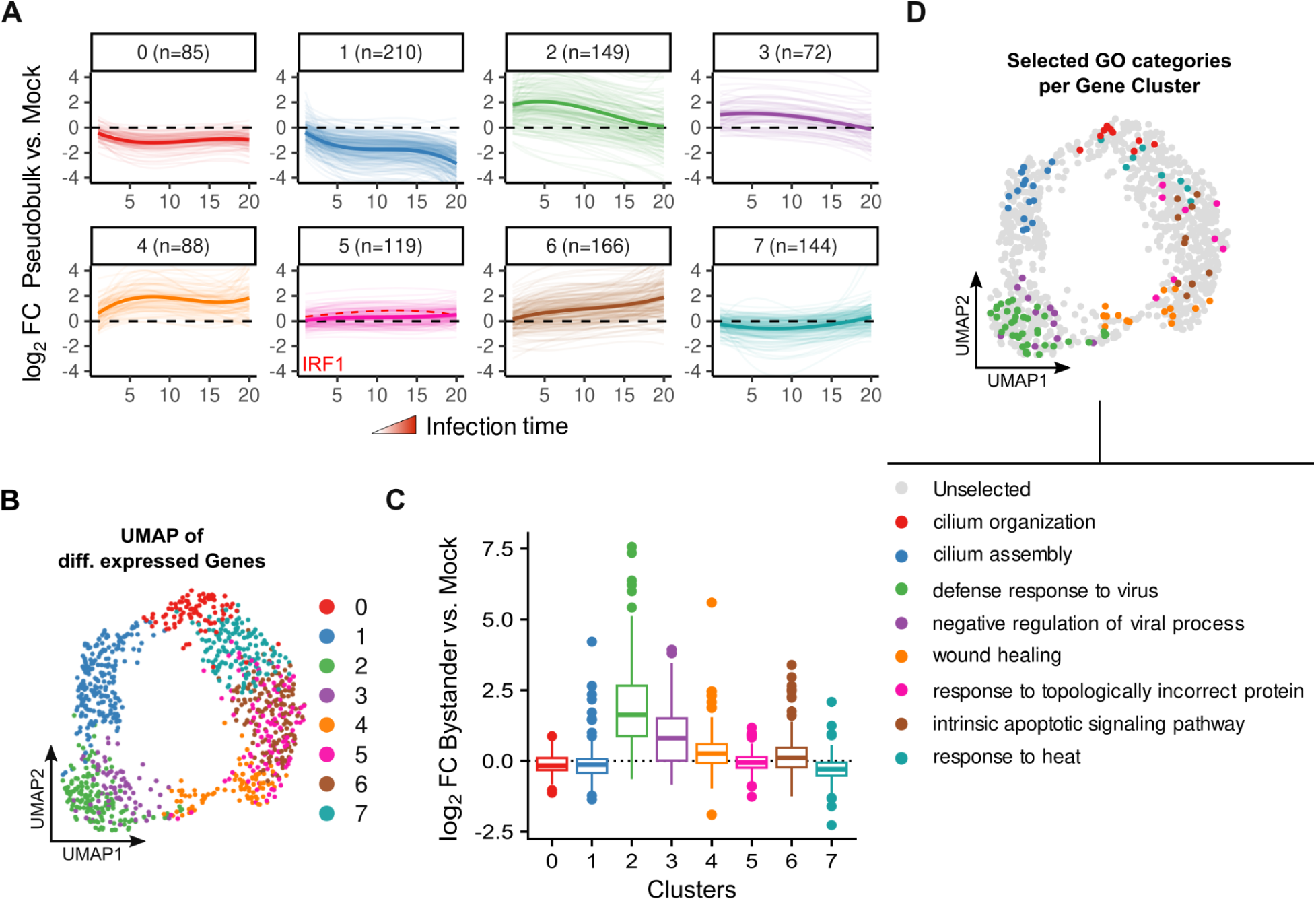
Uncovering viral load dependent transcriptional clusters by pseudo bulk analysis. (A) Mean LogFC values versus Mock per gene cluster and pseudobulk depict differing regulation patterns and magnitudes over infection time. All genes per cluster are shown as transparent lines, the opaque line represents the regression line over all genes. IRF1 is highlighted as a dashed red line. (B) UMAP representation of gene clusters of all differentially expressed genes over the Infection time, based on their LogFC values in all pseudobulks. (C) Boxplots showing the log2 fold changes of gene expression in bystander versus mock cells over all clusters. (D) UMAP representation of gene clusters. Genes associated with representative GO terms are highlighted.

In this gene UMAP visualization, the 8 clusters defined above together formed a continuous circle, with several gene clusters forming segments of the circle (clusters 0, 1, 4 and 7) or gene clusters forming the inner and outer parts of a given segment, respectively (clusters 2 and 3) (**Figure 7B**). Thus, this two-dimensional representation of host cellular gene expression faithfully captures the similarities and differences in cellular gene expression as viral load increases, thus visualizing how cells respond to and cope with increasing viral RNA load and tune the expression of specific gene sets. Of note, the gene UMAP also revealed that there was no group of genes that clearly segregated away from all other genes. Thus, these 8 clusters represent prototypical temporal/virus load-dependent profiles of genes, but there is indeed a continuum of temporal expression profiles and the gene UMAP describes how close a gene matches these prototypes.

The gene UMAP captures similarities and dissimilarities of genes in their temporal profiles irrespective of their behavior in bystander cells. Nevertheless, the regulation in bystander cells compared to mock was highly consistent with the initial infection time pseudobulks (**Figure 7C**). For instance, the most strongly upregulated genes in bystander cells were represented by genes in clusters 2 and 3. Notably, regulation in bystanders in all other clusters was less pronounced with clusters 0, 1 and 7 representing weakly downregulated genes in bystander vs mock cells and cluster 4 containing weakly upregulated genes. Genes in the remaining clusters 5 and 6 were unchanged. In summary, the expression in bystanders is consistent with expression at infection time 1 for all clusters suggesting that the bystanders represent the reservoir of cells that later initiate viral gene expression.

Notably, the highly upregulated genes in bystander cells were primarily interferon-stimulated genes confirming that both RSV-exposed bystander cells and RSV-infected cells mount a strong antiviral gene expression response that is dampened as viral load increases in the infected cells (**Figure 7A**, clusters 2 and 3). Interestingly, most other genes were regulated to a lesser extent suggesting that the IFN system was the major pathway that was activated upon RSV infection.

Several GO terms were overrepresented in distinct segments of the circle in the gene UMAP embedding in response to RSV infection, correlating with key biological processes (**Figure 7D**, **Supplementary Table 1**). Following genes around the circle, clusters 0 and 1 (jointly 29% of all regulated genes) consisted of genes continuously or progressively downregulated with increasing viral load, respectively. Both clusters were enriched with genes belonging to cilium assembly or organization, again confirming a rapid dedifferentiation of infected ciliated cells. Further cluster 1 genes (e.g., HLA-DPA1, HLA-DPB1, HLA-DMA) are involved in MHC class II complex assembly, implying RSV suppresses antigen presentation, potentially aiding immune evasion (**Supplementary Figure S8**).

Continuing on the circle, clusters 7 and 5 (∼25% of all regulated genes) showed minimal expression changes. Cluster 7 was dominated by protein folding genes (e.g., HSPA1A, HSPA6, HSPA8), while cluster 5 contained genes linked to starvation response, ER stress, and IFNγ signaling (e.g., IRF1, IFNGR2), highlighting a statistically significant but mild regulation of stress response proteins suggesting their importance in preserving cellular homeostasis.

Intermixed in the UMAP embedding with cluster 5 were cluster 6 genes (∼16% of all regulated genes), which were not regulated at the start of infection time, but steadily increased gene expression with viral RNA load. These genes reflected stress adaptation (e.g., DDIT4 inhibits mTORC1 to conserve resources) and apoptosis induction (e.g., CHAC1, DDIT4, CDA, IER3, BID). This suggests that increasing viral RNA load was accompanied by a broad cellular stress response ultimately leading to apoptosis when stress-dampening mechanisms are overwhelmed. To investigate this further, we treated ALI cultures with an inhibitor of apoptosis (zVAD-fmk), which increased the number of detectable RSV-infected cells (**Supplementary Fig. S6**). These results support the notion that with rising viral RNA load, RSV infection triggers apoptosis, maintaining the equilibrium between constant reinfection and cell death in our culture system.

Cluster 4 genes (9% of all regulated genes) were rapidly induced at low viral RNA loads and remained elevated. GO terms like “smooth muscle cell proliferation” and “wound healing” (e.g., TNFAIP3, CX3CL1, VEGFA, FGF2, CLDN1) suggested an early induction of tissue remodeling and epithelial barrier maintenance.

Finally, the next segment in the circle comprised cluster 2 and 3 genes (21% of all regulated genes), which were initially upregulated but showed a decline at higher viral RNA loads. Notably, cluster 2 formed a discernible bulge in the otherwise regular circle, and consisted of the most strongly regulated genes (e.g., *STAT1, CXCL10, TLR3*) that were linked to “defense response to virus,” highlighting interferon (IFN)-driven immune activation. Cluster 3 (e.g., *IFITM1-3, LY6E*) was associated with “negative regulation of viral process” featuring antiviral effectors that restrict viral replication. The strong induction and subsequent decline of these genes reinforced the role of type I/III IFNs in the immune response against RSV and viral counterregulation thereof.

Together, these clusters depict a dynamic progression from immune activation to cellular exhaustion, loss of epithelial integrity, and ultimately, apoptosis, emphasizing RSV’s ability to drive airway dysfunction.

### RSV fails to dampen IRF1 expression

Interestingly, not all ISGs are located in clusters 2 and 3 but instead, interferon regulatory factor 1 (IRF1) and Thioredoxin-interacting protein (TXNIP) (Cluster 5) show a progression that is less affected by increasing viral replication, possibly suggesting an inability of RSV to blunt their expression (**Figure 8A**). To further analyze the effect of IRF1 expression on RSV infection, we switched models and generated A549 cells stably overexpressing IRF1 (**Figure 8B-C**). Comparing RSV replication in vector control and IRF1 overexpressing A549 cells either by live-cell imaging (**Figure 8D-E**) or by flow cytometric analysis of RSV-infected cells (**Figure 8 F-G**), we observed a significant reduction of RSV infection in the cells with stable overexpression of IRF1 irrespective of the virus load used to inoculate the cells. As IRF1 is interferon-inducible we were interested in the effect of interferon treatment on RSV infection and replication. Therefore, we pretreated A549 vector control cells with two different doses of IFNα prior to infection with an RSV-A-GFP reporter virus at different MOIs and monitored RSV infection by live-cell microscopy (**Figure 8H-I**) and flow cytometry at 24h and 48h post infection (**Figure 8J-K**). As expected, pretreatment of cells with IFNα reduced RSV replication in a dose dependent manner. However, at later time points post infection, the same levels of RSV replication can be observed in IFNα treated compared to mock treated cells in case of the MOI1 and MOI 0.5 infections (**Figure 8H**), suggesting that at these inoculation doses RSV replication was delayed but not blunted, likely because RSV quickly prevents IFN-dependent antiviral effects. In contrast, viral replication in IRF1 overexpressing cells peaked at a lower level and never reached the same plateau as in the vector control cells irrespective of interferon treatment and viral inoculation dose (blue line in **Figure 8H**). To analyze whether interferon pretreatment had an additional inhibitory effect on RSV replication in IRF1 overexpressing cells, we treated IRF1 overexpressing A549 cells with the same doses as used in **Figure 8H-K** and monitored RSV replication by live-cell imaging (**Figure 8L-M**) as well as flow cytometry (**Figure 8N-O**). Given the fact that the viral replication in the IRF1 overexpressing cells is already weaker compared to the vector control cells (**Figure 8A-D**) it was interesting to see that the negative effect on viral replication can be even enhanced by treatment of cells with interferon, suggesting an additional mode of action independent from IFN signaling. As TXNIP is in the same cluster as IRF1 and shows a similar viral-load independent expression pattern, we performed the same experiments for thioredoxin-interacting protein (TXNIP). TXNIP has been described as “protein with many functions”, however one main function is regulating intracellular oxidative stress ^46^. In contrast to IRF1 overexpression, however, we were not able to see any significant changes in RSV replication upon TXNIP overexpression (**Supplementary Figure S9**).

**Figure 8:**
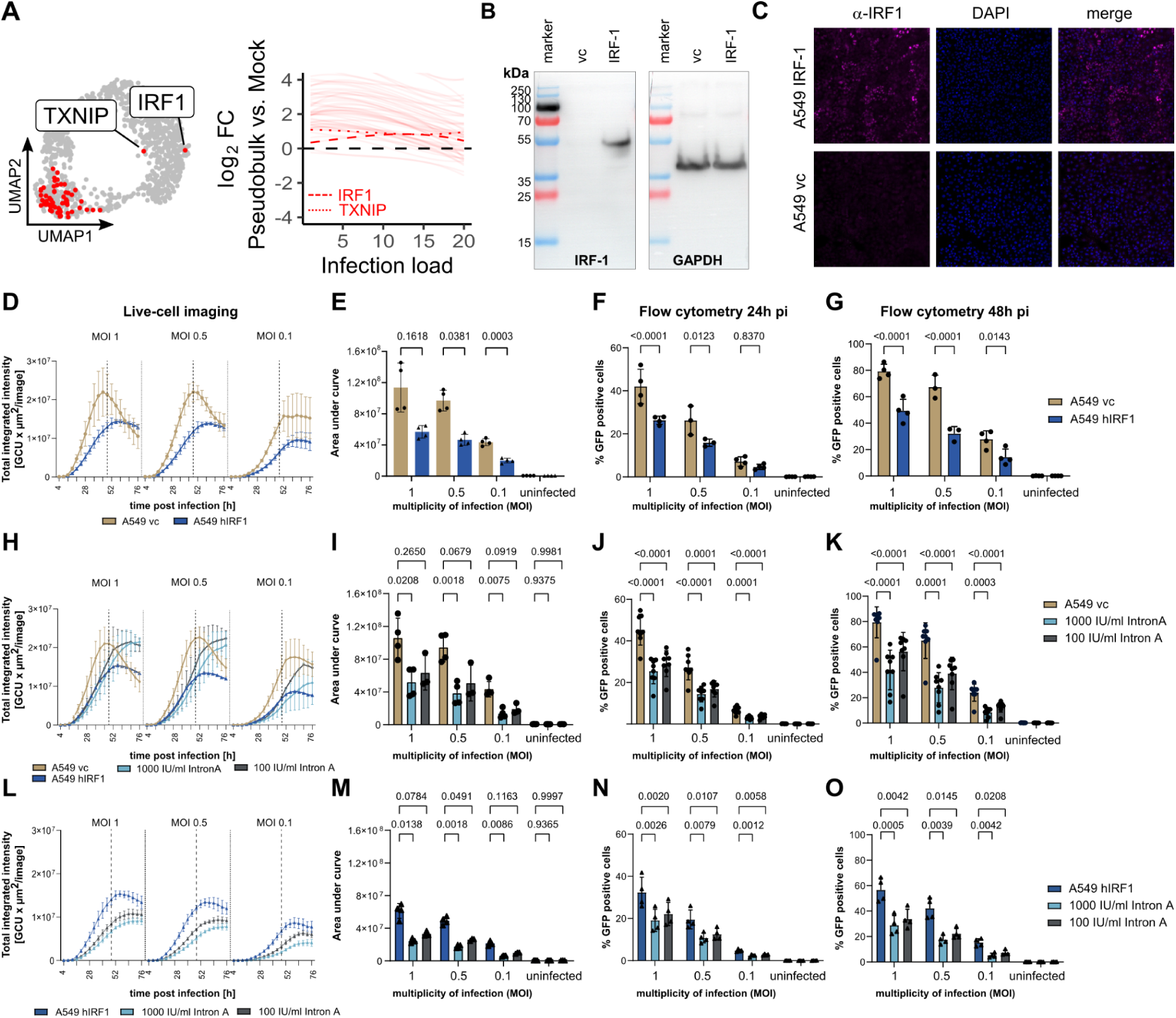
Influence of IRF1 overexpression on RSV infection. (A) UMAP representation of ISGs (red) across all clusters reveal differential regulation of IRF1 and TXNIP to all other ISGs (left). Mean LogFC values versus Mock per gene cluster and pseudobulk for ISGs depicts differing regulation patterns and magnitudes over infection time. IRF1 and TXNIP are shown as dashed and dotted lines, respectively (right). (B) Lysates from A549 cells either transduced with an empty vector control (vc) or with an IRF1 expression construct (IRF-1) were analyzed by Western blot analysis. GAPDH signal was used to show equal protein loading. (C) Immunofluorescence analysis using an IRF1 specific antibody (magenta, Proteintech #11335-1-AP) in combination with DAPI staining of vector control cells (A549 vc) and IRF1 overexpressing A549. One set of representative pictures is given. (D) A549 vector control (brown) or A549 cells stably overexpressing IRF1 (blue) were inoculated with an RSV-A-GFP reporter virus at indicated multiplicities of infection (MOI) for 3h. Viral inoculum was removed and live-cell imaging using an Incucyte machine started 4h post infection (pi) at a 10× magnification. Four pictures per well were taken every 4h for up to 76h pi. Total integrated green object intensity (GCU × µm^2^/image) was analyzed using the cell-by-cell software of Sartorius. (E) Area under the curve from (D) until the timepoint 48h pi was calculated and the statistical analysis was performed using a repeated-measurement one-way ANOVA in combination with Sídák’s multiple comparison test. Mean and std dev of n=4 independent experiments (D-E) as well as the results from the single experiments (symbols, E) is given. (F, G) Flow cytometric analysis of RSV-GFP positive cells at 24h (F) and 48h (G) post infection. Bars represent mean and std dev. of n=3-4 independent experiments. Additionally, the results from each independent experiment are depicted as symbol. Two-way ANOVA with Sídák’s multiple comparison test. (H-K) A549 vector control cells were pretreated for 16h prior to infection with an RSV-A-GFP reporter virus at indicated MOÍs with Interferon-2 alpha at two different concentrations. (H, I) Live-cell imaging and statistical analysis of n=3-4 independent experiments is given. For better comparison, the data from the A549 IRF1 overexpressing cells from (D, blue line) was added to panel (H). (J, K) Flow cytometric analysis of RSV-GFP positive cells 24h pi (J) and 48 pi (K) of n=8 independent experiments is depicted. Bars represent mean and standard deviation including the results for each independent experiment (symbol). Two-way ANOVA with Dunnett’s multiple comparison test. (L-O) A549 IRF1 overexpressing cells were treated with 2 different concentrations of interferon-2alpha 16h prior to infection with an RSV-A-GFP reporter virus at different concentrations and viral replication was monitored using live-cell imaging (L). (M) Area under the curve calculations until 48h post inoculation and statistical analysis of n=3-4 independent experiments using mixed-effects analysis and Sídáks multiple comparison test. (N, O) Flow cytometric analysis of RSV-GFP positive cells 24h pi (N) and 48h pi (O) from n=4 independent experiments is given.

## Discussion

The majority of studies investigating host responses to RSV infection have been carried out in cancer cell lines, so our knowledge of virus-induced host responses in primary human lung cells remains limited. A more comprehensive understanding of the effects of RSV on lung cells may provide important insights into the protective host response. In this study, we therefore analysed the kinetics and characteristics of host cell responses to RSV in an adult primary human pseudo-stratified airway epithelial cell model. The analysis focused on RSV and host transcriptional dynamics in bystander and infected cells, where cellular responses were analysed across the viral RNA load gradient, revealing continuous host transcriptional changes at the single cell level. We observed over 1,000-fold variation in viral RNA load between infected cells as early as 24 h post-inoculation, illustrating the enormous dynamics of viral replication and allowing us to stratify cellular responses across a wide range of viral loads. This work shows that RSV primarily, but not exclusively, infects ciliated cells. Some goblet and mucus-producing cells also showed signs of RSV infection. We show that a few of the infected ciliated cells are capable of responding with type I IFN production, thereby inducing a strong IFN response in the bystander cell population. However, in the infected ciliated cells, increasing viral load enhances the suppression of IFN induction and signalling, leads to cell dedifferentiation, loss of MHC class II gene expression, upregulation of the cellular stress response and ultimately apoptosis. Finally, we found that mRNAs of selected ISGs such as IRF1, a transcription factor that regulates many genes, are not attenuated by RSV and that increased IRF1 expression may be a way to protect cells from RSV infection and pathogenesis.

Consistent with previous reports ^30–32^, RSV mRNA expression decreased from the 3’ to the 5’ end of the viral genome and G protein mRNA was over abundant relative to its position in the viral genome ^47^. Surprisingly, however, we also observed an overabundance of SH and M2 mRNAs, which to our knowledge has not been described before. Several mechanisms have been proposed to explain deviations from the RSV transcriptional gradient. A contributing factor could be differential transcript stability. Alternatively, variable transcription initiation probability at the gene start sequences or polymerase recycling, i.e. repeated transcription of the same gene before proceeding downstream, can amplify mRNA output for specific genes such as G, SH or M2. While we cannot fully rule out that our results are partly due to the estimation of mRNA abundance based on single cell mRNA sequencing results or the use of an RSV reporter virus, these data nevertheless suggest that transcriptional regulation in RSV may be more complex than previously thought, thus fine-tuning gene product abundance.

Several lines of evidence show that RSV primarily, if not exclusively, infects ciliated lung epithelial cells ^9–12^. In well-differentiated primary airway cultures, RSV infection is mostly found in ciliated cells, which form the apical layer of the pseudostratified epithelium. By contrast, there is no direct evidence that mucus-secreting goblet cells (which intersperse between ciliated cells in the airway epithelium) are infected by RSV and produce viral RNA, reporter genes or other RSV antigens. Exceptions have been noted, when the epithelium is breached under experimental conditions so that RSV also infected basal cells ^16^. This ciliated-cell tropism has also been observed *in vivo*: Histopathologic studies of infected tissue reveal RSV antigen in columnar airway epithelium of large airways and in bronchiolar ciliated and non-ciliated cells ^13,48,49,50^. In one study, the epithelia of small bronchioles were peripherally infected, basal cells were not found to be infected, but type 1 and 2 alveolar pneumocytes were also infected ^13^. In our analysis, some goblet and mucus-producing cells also showed signs of RSV infection including GFP expression and virus RNA, albeit only with low RSV copy numbers (**Figure 3A**). Interestingly, goblet/mucus marker gene expression of these RSV-infected goblet- and mucus-like cells was slightly skewed compared to the mock-treated goblet and mucus cell populations. This suggests that RSV infection may also cause dedifferentiation of these cell types. We did not find any goblet- or mucus-like cells with a high viral load. This may suggest a lack of host factors or strong antiviral defenses in these cells preventing further increases in viral load and protecting these cells. However, we cannot rule out the possibility that these cells die at a low viral load, or that massive transcriptional reprogramming due to high viral load has led to their labelling as basal cells.

Our analysis of cellular changes upon RSV infection revealed a significant loss of basal cells in the bystander cell population. At the same time, most infected cells shared gene expression characteristics of both ciliated and basal cell populations, with low virus load cells were more similar to ciliated cells and high virus load cells shared stronger similarity with basal cells. Profiling of cellular marker genes showed that with increasing virus load, RSV-infected cells de-differentiate, lose ciliated cell characteristics and adopt a basal cell-like gene expression profile. In keeping with previous reports, we conclude that RSV primary infects ciliated cells. These cells are unable to arrest infection and consequently progress to a high viral load, leading to dedifferentiation and ultimately cell death. To further investigate cellular processes occurring along the increasing viral load in ciliated cells, and to potentially identify pathways that could be targeted to halt infection, we performed a more comprehensive analysis of transcriptional changes across the viral load. The most notable findings from this analysis are a virus load-dependent decrease of type I IFN induction and signalling, reduced expression of MHC class II subunit transcripts, and an increase of cellular stress responses that ultimately lead to apoptotic cell death. However, we also found that selected interferon-stimulated genes (ISGs) such as interferon regulatory factor 1 (IRF1), a potent transcription factor that controls numerous host antiviral genes, were not downregulated by RSV infection. Furthermore, we showed that IRF1 overexpression, in contrast to exogenous type I IFN, significantly reduced RSV infection, suggesting its potential as a prophylactic and therapeutic intervention for RSV infection. Finally, we show that cellular dedifferentiation is selective for RSV-infected cells and does not involve bystander cells. This demonstrates that cell dedifferentiation is primarily an RSV-driven process. Bystander cells, which include numerous uninfected ciliated cells exhibited a robust ISG response already at day 1 post infection and remained largely unaltered otherwise compared to mock cells. Their antiviral state due to stable ISG expression indicates that they are protected from infection or the onset of viral gene expression. Nevertheless, the maintenance of low viral RNA load cells also at days 3, 5 and 7 suggests continuous reinfection events throughout the seven days of the experiment. The depletion of basal cells in the cultures exposed to RSV is consistent with a model where basal cells constantly differentiate to replenish the ciliated cell compartment, and that a subset of the freshly differentiated ciliated cells are then successfully infected by RSV.

Our work confirms previous studies showing that RSV induces a type I and III IFN response ^36–43^. However, we extend this work by showing which cell types produce type I and III IFNs and by quantifying the efficacy of IFN signaling in bystander cells and in relation to virus load in the infected cells. Our results show a pronounced virus load-dependent silencing of IFN responses, induction of type I and III IFN only in a fraction of infected cells and a strong IFN response in all bystander cells. Thus, most RSV-infected cells are unable to produce IFN. This may explain why some children have undetectable IFN levels in their respiratory swabs and why only low levels of IFNs are detectable when human primary lung cells are infected by RSV ^36,38–40^. However, this confined induction of type I and III IFNs is apparently sufficient to keep RSV from spreading through the ALI culture. Indeed, blunting IFN signalling by pharmacological means increased RSV infection suggesting that IFN indeed is antiviral. However, exogenous addition of IFNα did not downregulate RSV infection. This indicates that IFN is fully operational and that the main mechanism of NS1/NS2 is exerted downstream of IFNAR. The early burst of the ISG response on day 1 in both bystander and infected cells could be due to carry-over of residual IFN from the producer cells, or to early unattenuated IFN production and signalling prior to accumulation of viral antagonists or cellular silencing of excessive IFN signalling. In this latter scenario the continuous IFN stimulation may downregulate ISG induction on days 3, 5 and 7.

It is unclear why only a few infected cells produce type I and III IFN. This could be because RSV infection is slower to ramp up in these cells, possibly due to fewer infecting viruses or other constraints, such as the levels of cellular host factors that drive transcription or replication. In such a scenario, the slow accumulation of viral antagonists could allow cells to signal type I and III IFNs before effective blunting occurs. Alternatively, higher endogenous expression of pattern recognition receptors and/or key transcription factors may allow some cells to respond to rapidly increasing viral loads (Figure 6).

Human studies provide conflicting evidence regarding the correlation between type I and type III IFN levels in nasopharyngeal secretions and the severity of RSV disease in infants, with one study reporting an inverse correlation and another a direct correlation ^38,51^. These discrepancies may be due to differences in sampling time relative to the course of infection and/or sampling procedures. On the one hand, we observed that (I) RSV is unable to spread through the entire potentially susceptible population of ciliated lung epithelial cells over seven days of *in vitro* culture (II) and that blunting of IFN signalling enhances virus spread. In addition, relatively few cells were infected by RSV (0.7 to 3.2%), suggesting that IFN-dependent antiviral responses provide a strong barrier to RSV infection. On the other hand, exogenous addition of IFN was unable to suppress RSV infection in this model over a seven-day time course. Taken together, this suggests that type I and type III IFN signalling from a few infected cells is necessary and sufficient to keep RSV infection in check at steady state levels, but not sufficient to clear the infection. Given the blunting of type I and III IFN production at high viral load classes and of IFN signalling in infected cells with high viral load, it is tempting to speculate that the kinetics of IFN induction may be an important set point that co-determines the course and outcome of RSV infection, both at the single cell level and in infected individuals. For example, high-dose transmission of multiple virus particles could favour the rapid establishment of high viral loads, the accumulation of viral antagonists and the establishment of a broad cellular repertoire that seeds deep lung infection and prolonged inflammation. Alternatively, higher baseline expression of pattern recognition receptors such as RIG-I and MDA5 could favour rapid induction of type I and III IFNs and milder courses of infection. In this context, it will be interesting to investigate the mechanisms that allow some RSV-infected cells to produce IFNs while the majority of infected cells fail to induce these cytokines.

The observation that RSV is not eliminated from the culture but also does not spread to all available ciliated cells suggests that the recruitment and function of innate and adaptive immune cells is critical for clearance of RSV infection. This finding is also consistent with *in vivo* observations in immunosuppressed patients. In such patients, RSV infection can persist for weeks without viral clearance and is associated with increased severity and mortality ^52,53^. Our *ex vivo* analysis suggests that sufficient new susceptible ciliated cells are available over the course of 7 days, probably due to differentiation from basal cells and resistance to infection due to IFN-triggered antiviral defenses. Considering the essential absence of differentiating and proliferating basal cell compartments in our RSV-exposed culture, we suggest that RSV infection demands maximum regeneration of the system to replenish ciliated cells that are continuously depleted by RSV infection. It is conceivable that, depending on the number of infected cells and the site of infection *in vivo*, RSV infection could outcompete the regenerative capacity *in vivo*, leading to fatal disease in immunosuppressed patients. In light of these findings, therapies that block the activity of NS1 and NS2 function may help to limit RSV infection in patients with an ineffective adaptive immune response. Alternatively, boosting endogenous defence may be a strategy to alter the balance between RSV replication and cellular clearance. With this in mind, we searched for potentially antiviral cellular responses that are not effectively blunted by RSV during infection. One interesting factor in this regard is IRF1. We show that, at least at the mRNA level, RSV infection is unable to dampen the expression of this transcription factor.

IRF1 is known to inhibit the replication of several viruses by partially interrelated mechanisms. On the one hand, IRF1 has been shown to prevent the interaction between IRF3 and protein phosphatase 2A (PP2A), which prevents the dephosphorylation of IRF3 and thus the silencing of its activity ^54^. As a result, IRF1 is maintained in its activated state, which enhances type I IFN production and stimulation of ISGs. On the other hand, IRF1 is able to maintain the expression of certain antiviral ISGs, such as OAS2, BST2 and RNASEL, independently of IFN signalling. These and other IRF1-dependent genes could establish an antiviral state in lung epithelial cells, thereby reducing their susceptibility to respiratory viruses ^55^. It also facilitates histone H3K4 monomethylation (H3K4me1) at promoter-enhancer regions of ISGs, priming them for rapid expression upon viral detection. It also recruits BRD4, a transcriptional coactivator, to enhance IFNβ and IFNλ expression following pathogen recognition receptor (PRR) stimulation. Finally, IRF1 regulates the constitutive expression of PRRs such as TLR2 and TLR3, thereby increasing cellular sensitivity to viral nucleic acids. This amplifies the downstream activation of IRF3 and NF-κB, creating a positive feedback loop for antiviral responses.

Importantly, when we ectopically overexpressed IRF1 in A549 cells, we observed a clear antiviral effect against RSV. In fact, this antiviral effect was not overwhelmed by high-dose RSV challenge, unlike the effect of, for example, IFN pre-treatment, which was blunted by high viral inoculum. We also found that ectopic expression of IRF1 has an additional antiviral effect to IFN treatment. Thus, in principle, these therapeutic approaches appear to work together to silence RSV infection. Of course, these latter therapeutic approaches have only been investigated in a human lung cancer cell line and using a constitutive overexpression system. Therefore, confirmation in more authentic lung cell models and ultimately *in vivo* is important. It should also be noted that IRF1 is a transcription factor that is only expressed in cells where the transgene is delivered to. Thus, unlike IFN, which is secreted and acts both auto- and paracrine, IRF1 is expected to protect only those cells that take up the transgene.

In terms of our single cell transcriptional profiling, our model is reductionist in that it does not and cannot take into account the role of resident and infiltrating immune cells. While this is a limitation that prevents the measurement of the influence of cytokine networks and innate effector functions of immune cells and their secreted factors on the dynamics of RSV infection and pathogenesis *ex vivo*, we consider this focus also a strength of our study. We used six independent human adult donors to exclude artefacts due to inherent lung diseases of individual patients. We also confirmed that the main findings were reproduced in each donor. Finally, we only had access to adult human lung tissue. Therefore, it is currently unclear whether and to what extent our findings can be extended to RSV-host cell interaction in human paediatric lung cells.

This study opens up several interesting new perspectives. First, it will be interesting to investigate why RSV infection appears to be abortive in goblet and mucosal cells. It is possible that this is related to a specific antiviral control programme of these cells that is inactive or less active in ciliated cells. Secondly, it will be of interest to investigate the mechanisms that allow some of the infected ciliated cells to mount type I IFN responses whereas others fail to do so. Understanding the underlying principles may provide a means to enhance endogenous defence in a wide range of lung cells, thus providing a greater barrier to RSV infection and ultimately increased resilience to RSV. Finally, the observation that RSV is unable to attenuate IRF1 transcription and that, at least in A549 cells, there is a significant prophylactic benefit of IRF1 overexpression provides a new basis for testing IRF1 in RSV (and possibly other viruses) prophylaxis.

## Methods

### Ethic statement

Adult patients gave informed consent for the donation of lung samples and all steps were in compliance with good clinical and ethical practice and approved by the local ethical committee at Hannover Medical School (permission number 3346/2016).

### Cell lines, ALI culture and virus

A549 cells (ATCC; CCL-185) were cultured in F12K NutMix media supplemented with 10% heat-inactivated FCS (Capricorn Scientific), 1% NEAA (Gibco), 2 mM L-glutamine (Gibco), 100 U/ml Penicillin and 100 U/mL Streptomycin (Gibco) at 37°C and 5% CO2 in a humidified incubator. To generate cell lines stably expressing hIRF1 or TXNIP, A549 cells were either transduced with lentiviral particles encoding the vector control (pWPI-BLR) or a lentiviral vector encoding for human IRF1 (pWPI-hIRF1-BLR) or human TXNIP (pWPI-TXNIP-BLR), respectively. Three days post transduction, cells were selected for efficiently transduced cells by addition of blasticidin as selection antibiotic.

Primary human airway epithelial cells cultured under air-liquid-interface (ALI) conditions were cultivated and differentiated as described elsewhere ^56^. In brief, human airway epithelial cells were isolated from the bronchus of explanted human lungs by enzymatic digestion for 48h at 4°C. Cells were seeded on collagen I/III-coated flasks for proliferation prior to seeding on collagen IV-coated transwells (0.4 µm pore size, polyester membrane inserts, Corning Costar). All cells were routinely tested negative for mycoplasma contamination (Eurofins MWG).

Recombinant reporter virus rHRSV-A-GFP was described elsewhere ^33^. Virus stocks were prepared in HEp-2 cells (ATCC, CCL-23). For harvesting, infected cells were scraped and vigorously vortexed to release cell-bound virus particles. Virus preparation was centrifuged for 10 min at 1000 × g to remove cell debris, supplemented with virus stabilizer (final concentration 100 mM MgSO_4_, 50 mM HEPES, pH 7.5), aliquoted and snap frozen in liquid nitrogen. Virus titer was determined using the limiting dilution assay (TCID_50_).

### Cell preparation and library preparation for single-cell RNA-seq

Differentiated primary human airway epithelial cells from six different donors were inoculated with recombinant RSV-A-GFP reporter virus for 2h at 37°C (1.4×10^5^ infectious virus particles/transwell) or with conditioned media generated in parallel to the virus stock. After 2h, the viral inoculum was removed and the apical compartment was washed twice with Hank’s balanced salt solution (HBSS). At different timepoints post virus inoculation, cells were washed apically with HBSS prior to cell detachment by addition of trypsin/EDTA and incubation at 37°C. Then, cells from the different donors belonging to the same time point were pooled prior to hashtagging (TotalSeq-A Antibodies, BioLegend). RSV-infected (GFP-positive) cells were sorted using a FACS Aria III and 3,000 GFP-positive and 6,000 bystander cells per time point were loaded in the 10x Genomics controller. In parallel and simultaneously, non-infected cells were trypsinized, hash-tagged for the time point and applied for the 10x Genomics (see below).

Cell suspensions at a density of 400 cells/µl in PBS+0.04% BSA were prepared for single-cell sequencing. Chromium™ Controller was used for partitioning single cells into nanoliter-scale Gel Bead-In-Emulsions (GEMs). Briefly, 5-10,000 cells per reaction were loaded for GEM (gel bead-in-emulsion) generation and barcoding. Libraries were constructed using the Chromium Single Cell 3′ Reagent Kit v2 and v3 (10x Genomics), following manufacturer’s instructions. A SimpliAmp Thermal Cycler was used for amplification and incubation steps (Applied Biosystems). Libraries were quantified by a QubitTM 3.0 fluorometer (Thermo Fisher Scientific) and quality was checked using a 2100 Bioanalyser with High Sensitivity DNA kit (Agilent Technologies). Samples were pooled and sequenced using the Illumina NovaSeq 6000 in a paired-end mode: Read1, 26 cycles; Index1, 8 cycles; Index2, 0 cycles; and Read2, 98 cycles. CellRanger v2.1 or later software was used to process the raw FASTQ files, aligned the sequencing reads to GRCh38 genome built and generated a filtered unique molecular identifier (UMI) expression profile for each cell.

### Live-cell imaging

Cells were seeded into 96-well plates at 7×10^3^ cells/well in a total of 100 µl F12K NutMix cplt media. Eight hours post infection, cells were treated with indicated concentrations of human interferon alpha-2b (IntronA, MSD Sharp&Dome GmbH) where indicated. 16h post IFN treatment, cells were inoculated with an RSV-A-GFP reporter virus at indicated multiplicity of infection (MOI) for 3h in a total of 50 µl. After inoculation, viral inoculum was removed and replaced by 100 µl media supplemented with IFN as indicated. Scanning of cells started 4h post inoculation using an Incucyte SX5 machine (Sartorius BioAnalytical Instruments). Four pictures each from at least 2 wells were scanned every 4h using a 10× magnification. Total green integrated intensity (GCU× µm^2^/image) was accessed using the Incucyte software (GUI Version 2023A, Controller Version 2023A Rev1, Firmware Version 20231.1.4.0 RTM).

### Flow cytometry analysis

Cells were seeded into 96-well plates at 1×10^4^ cells/well the day prior to infection with the recombinant reporter virus RSV-A-GFP at different multiplicities of infection for 4h. 24h or 48h post inoculation, cells were trypsinized and fixed in 3% paraformaldehyde in PBS for at least 30 minutes. Cells were stored at 4°C upon flow cytometric analysis using a SONY spectral analyzer SA3800 and results were analyzed using FlowJo V10. Gating strategy is depicted in Supplementary Figure S1C.

### Immunofluorescence analysis

RSV-A-GFP-infected differentiated primary human airway epithelial cells grown under ALI conditions were fixed with 4% formalin solution for at least 30 min at room temperature from the apical and basolateral side. After removal of the fixation solution, both compartments were washed three times with PBS prior to addition of the blocking buffer (50 mM NH_4_Cl, 0.1% saponin, 2% BSA in PBS, pH 7.4) for 1h. Blocking buffer was replaced by antibody dilutions to the apical compartment and incubated for 2h at room temperature. After binding of 1^st^ antibody, the apical surface was washed three times with blocking buffer prior to addition of the conjugated secondary antibodies to the apical compartment for 1h. Unbound antibodies were washed away by 3× PBS washes, the membrane was excised from the transwell and mounted on glass cover slides. Immunofluorescence acquisition and analysis were performed using an Olympus microscope (FV3000 and CellSens software, Olympus).

| **Antibody<sup>57</sup>** | **Supplier** | **Catalogue** | **Lot.** |
| --- | --- | --- | --- |
| CD49f (Integrin alpha 6),<br>Clone eBioGoH3 (GoH3) | Invitrogen | 14-0495-82 | 2518417 |
| β-Tubulin-Cy3,<br>Clone TUB2.1 | Sigma | C4585 | 088M4821V |
| Cytokeratin 5 [EP1601Y] | Abcam | ab52635 | GR3292032-15 |
| ARL13B | Proteintech | 17711-1-AP | 00103765 |
| Mucin 5AC | Santa Cruz | sc-21701 | G3117 |
| CD271-PE | BioLegend | 345106 | B360532 |
| AF647 anti-rat | Invitrogen | A21247 | 1750833 |
| AF647 anti-rabbit | Invitrogen | A21244 | 1741783 |
| AF568 anti-rabbit | Invitrogen | A10042 | 1891789 |
| AF647 anti-mouse | Invitrogen | A21235 | 2284596 |

### Statistical analyses

Statistics of live-cell microscopy was analyzed using the area under the curve (AUC) method. Therefore, AUC was analyzed using GraphPad prism (Version 10.0.2) for each independent experiment. RM one-way ANOVA with Sidák’s correction for multiple comparisons was used. Error-bars and replicate numbers are defined in the respective figure legends.

### scRNA-seq data processing, quality control, cell type annotation and general data analysis

The fastq-files were mapped using the cellranger pipeline (version 7.0.0). The human genome sequence was taken from the ENSEMBL database (version 90) and the RSV sequence was taken from Genbank (Accession number MK816924). Data analysis was performed in R (version 4.3). Filtering, normalization, feature selection and dimensionality reduction were performed using the Seurat package (version 4.3.0). Visualization was done using the ggplot2 package (v. 3.5.1).

At first, time points were demultiplexed using the Seurat package. First, the hashtag read counts were normalized using NormalizeData with the centered log ratio transformation (CLR) as normalization method and then, the HTODemux function was used with positive.quantile = 0.998. Subsequently, all hashtag negative (n=7238) and doublet cells (n=849) were removed.

We kept all features that were detected in more than 3 cells and filtered cells according to the following statistics: Number of features (1,500< x < 10,000), number of UMIs (5,000 < x < 100,000), percentage of mitochondrial reads (< 30%), number of viral UMIs (<50 in bystander cells, >100 in infected cells, see **Fig. 1B, Suppl. Fig. 2A**).

To annotate the cell types, we used the IntegrateData function of the Seurat package (v. 4.3.0) with standard parameters to integrate all mock cells of our data into a single cell atlas of the human lung ^34^ and calculated the nearest neighbor graph (k=50) on the integrated data set. Subsequently, we iterated over all unannotated mock cells and labeled each cell according to the most prevalent cell type in its nearest neighborhood. After mock cell annotation, we integrated the mock and unannotated virus-exposed cells using standard parameters, calculated the nearest neighbor graph (k=20) and labeled the virus-exposed cells according to their neighborhood to the mock cells as described above.

We removed all cell types with less than 10 cells and further scrutinized all remaining cells after clustering. This led us to remove another set of cells which survived the general filtering thresholds defined above but had more than twice the number of mitochondrial reads and an order of magnitude less detected UMIs than all other cells (see **Suppl. Fig. 1B-D**).

Dimensionality reduction was performed on the integrated assay using the RunPCA function with standard parameters and the RunUMAP function with 35 dimensions from Seurat. The viral RNA load of cells was calculated as the log of total sum of normalized viral gene counts per cell.

The donor cells were demultiplexed by genetic variants using souporcell (v. 2.0) from a singularity container (v. 3.9.1+84-g510833307) with the following command: “singularity exec souporcell.sif souporcell_pipeline.py -i bamfile.bam -b barcodes.tsv -f reference.fasta” with the bam-file and barcodes from the cellranger output and the human genome sequence from the ENSEMBL database (version 90). We discarded all cells that were not assigned as singlets.

### Viral gradient plot and manual RSV annotation

For the viral gradient plot (**Fig. 1E**), we inspected read coverage in the RSV genome in a genome browser (Display function of the GEDI toolkit - version 1.0.6). We created a manual annotation of the eleven poly-A sites in the RSV genome by extending the visually identified read count peaks identified in the genome browser by 200 bp on both sides. We then reran the cellranger pipeline with the updated annotation and then determined the proportion of raw read counts for each viral gene relative to the total viral read counts per cell and plotted them in order based on their position in the viral genome. The manual annotation of the RSV genome for 3’ end peaks is available as **Supplementary Data 1**.

### Gene ontology analysis

Gene ontology analysis was performed in R using the clusterProfiler package (version 4.8.1) using all genes in the respective cluster as the targets, all genes in the dataset as the background and the org.Hs.eg.db package (version 3.17.0) as the database. We adjusted p-values using Benjamini-Hochberg correction and set a q-value threshold of 0.05 for all GO analyses and a logFC threshold of −0.5 for the GO analysis of downregulated genes in basal-like compared to ciliated cells (**Fig. 3D**)

### Infection time analysis of infected cells

For the infection time analysis we only included cells annotated as ciliated, proximal ciliated, proximal basal and basal cells. We sorted the infected cells by the number of viral reads and binned them into 20 pseudobulks each consisting of n=85-86 cells, pooling reads of all cells per pseudobulk. Similarly, we pooled the reads of all mock and bystanders into a pseudobulk, respectively. We then calculated normalized log fold-changes versus mock cells using the PsiLFC function of the lfc package (version 0.2.3) with standard parameters ^58^. Differential gene expression analysis between pseudobulks of infected cells on the 5,000 most variable genes was performed using the DESeq2 package (version 1.40.1). We included all genes with an adjusted p-value < 0.01 (n=1033) in the analysis.

To cluster genes by their progression over infection time rather than absolute log fold change (LFC) values, we centered each gene by subtracting its mean LFC from all LFCs of that gene. A Seurat object was created from the LFC values of the bystander and infected pseudobulks, loading genes as observations and pseudobulks as features. The data was then scaled and PCA was conducted using the ScaleData and RunPCA functions with standard parameters and the nearest neighbor matrix (using the FindNeighbors function and the first 20 dimensions) was calculated. Finally, clustering was performed using the FindClusters function with a resolution of 1.5. The gene-based UMAP was then constructed with the RunUMAP function on the first 20 principal components. In all cases, euclidean distance was used.

The line plots (**Fig. 7A**) show the smoothed LFCs vs. mock of all genes in all pseudobulks per cluster and a regression line over all genes per cluster. The lines were created using the geom_smooth function of ggplot2 with “lm” as the smoothing function and the formula y ∼ splines::bs(x, 3). To improve the visualization and remove outliers, the y-axis was limited to LFCs −4 < x < 4.

## Data availability

Data and code to reproduce all results are available at Zenodo (https://doi.org/10.5281/zenodo.15705874) and data can be browsed under http://wvir052.virologie.uni-wuerzburg.de:3839/6780754e77dccbdaef47451a4f105a04/

## Acknowledgements

Funded by the Deutsche Forschungsgemeinschaft (DFG, German Research Foundation) under Germanýs Excellence Strategy-EXC 2155-project number 390874280 to TP, under SFB 1583/1 – Project number: 492620490 (Z02) to AES and FE, and under ER 927/2-1 to FE. TP received funding from the Volkswagen Stiftung within the Niedersächsische Vorab initiative “COALITION-Communities allied in infection” and the “INDIRA: INtegrative Data analytIcs for Respiratory syncytial virus risk Assessment” projects. BW was funded by the German Research Foundation (DFGSPP2014: 347346497, 447746988 and 347368182, KFO311: 286251789), the German Centre for Lung Research (DZL): BREATH (Biomedical Research in End Stage and Obstructive Lung Disease Hannover; DZL: 82DZL00201). BW is a member of the DFG-SPP2014, Hannover, Germany. We are grateful to Marie-Anne Rameix-Welti and Jean-François Eléouët for the kind gift of the RSV reporter viruses and to Ronald Dijkman for inspiring discussion of the single-cell data. We acknowledge the use of large language models (LLMs) to assist in the formulation and optimization of text in this manuscript, particularly for improving overall language and refining the introduction section. All content generated with LLM support was reviewed for accuracy, and the authors take full responsibility for the final text.

## Supplementary Figures

**Supplementary Figure S1:**
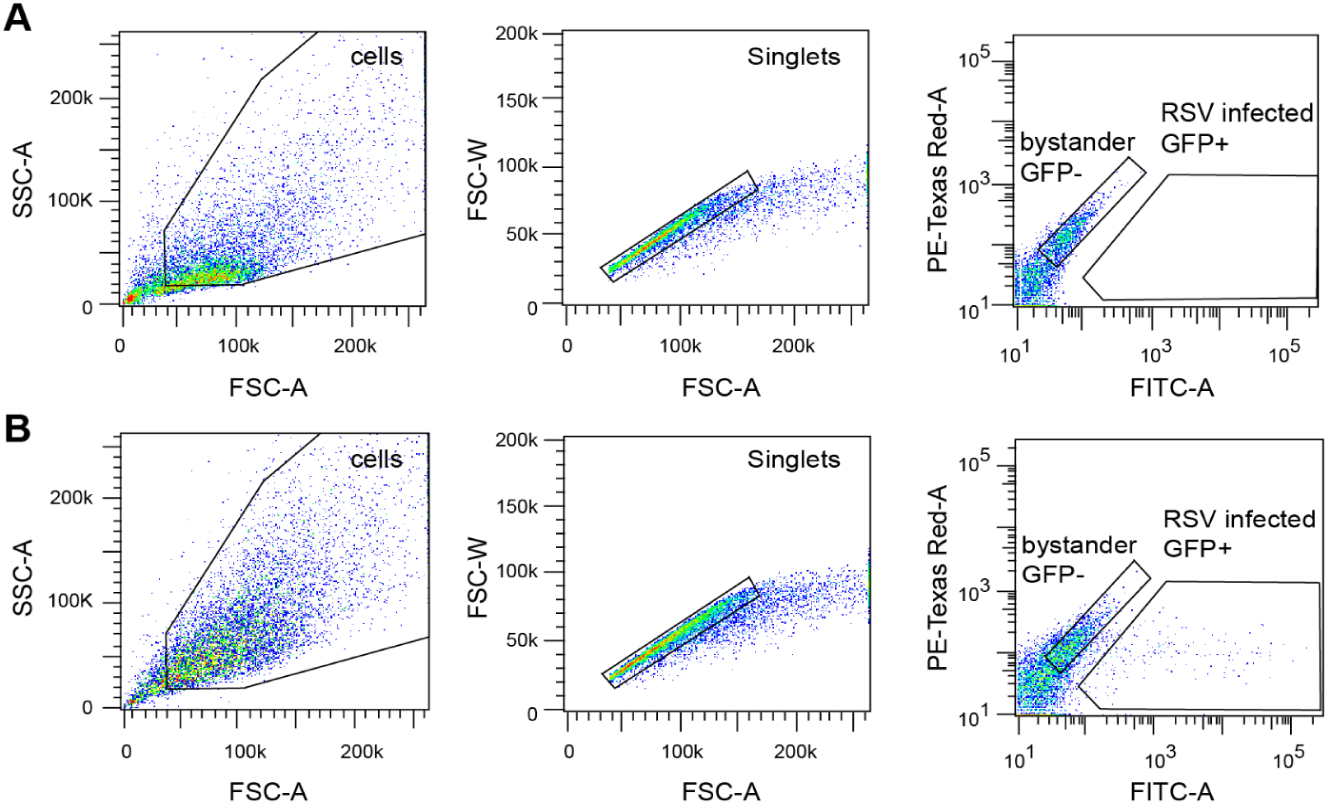
Representative of FACS gating strategy. **(A)** Uninfected and **(B)** RSV-GFP infected primary human airway epithelial cells 3 days post inoculation were FACS sorted using a BD FACSDiva system. Cells were gated for living and singlet cells prior to sorting for GFP-negative bystander cells and GFP-positive RSV-infected cells. (C) Gating strategy for A549 cell lines infected with RSV-A-GFP (MOI1) for 24h.

**Supplementary Figure S2:**
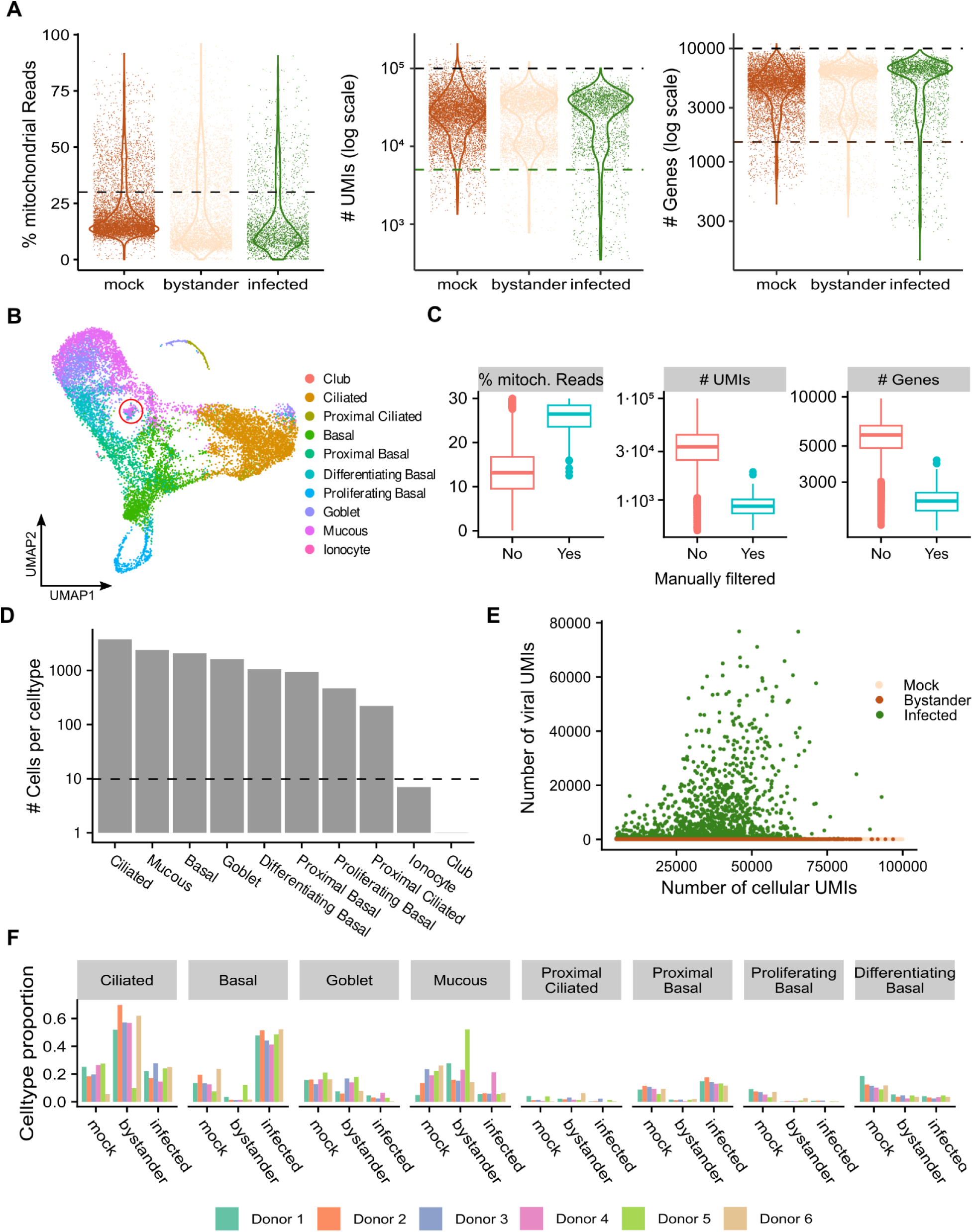
Data quality and processing of single-cell RNA-seq data. **A** Violin plots of the percentage of mitochondrial reads (left), number of detected unique molecular identifier (UMIs) (middle) and number of detected genes (right) per cell, split by infection state. Dashed lines indicate the filtering cut-offs. **B** UMAP representation as in Figure 1D of annotated cells that passed basic quality control measures (12537 cells). Ionocyte (7 cells) and club cells (1 cell) as well as cells with a combination of very high mitochondrial reads and low read and feature counts (198 cells, red circle) were removed from the analysis. **C** Boxplots of the percentage of mitochondrial reads, number of detected UMIs and number of detected genes per cell for the manually removed cells (red circle in **A**) and the remaining data set. **D** UMAP representation of single-cell RNA-seq annotated and color-coded according to lung atlas-based cell type annotation. **E** Scatter plot showing the number of cellular UMIs versus the number of viral UMIs in mock (beige), bystander (brown) and infected (green) cells. **F** Relative proportion of cell profiles per infection state decomposed according the donor of origin.

**Supplementary Figure S3:**
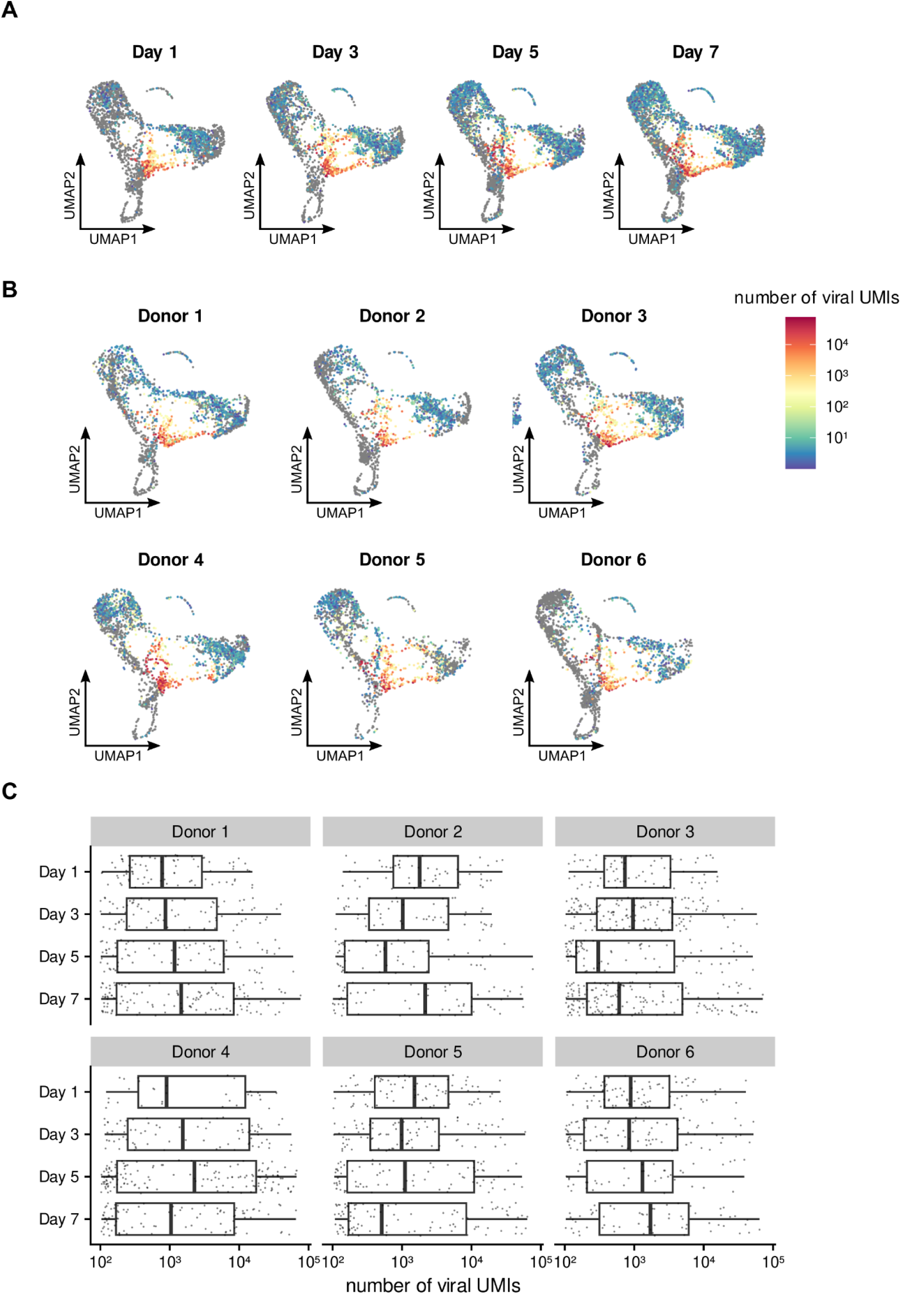
Distribution of viral UMIs across time points and donors. (A) UMAP representation of viral UMIs in infected cells across all time points. (B) UMAP representation of viral UMIs in infected cells across all donors. (C) Boxplots showing the number of viral UMIs in infected cells across all time points and donors.

**Supplementary Figure S4:**
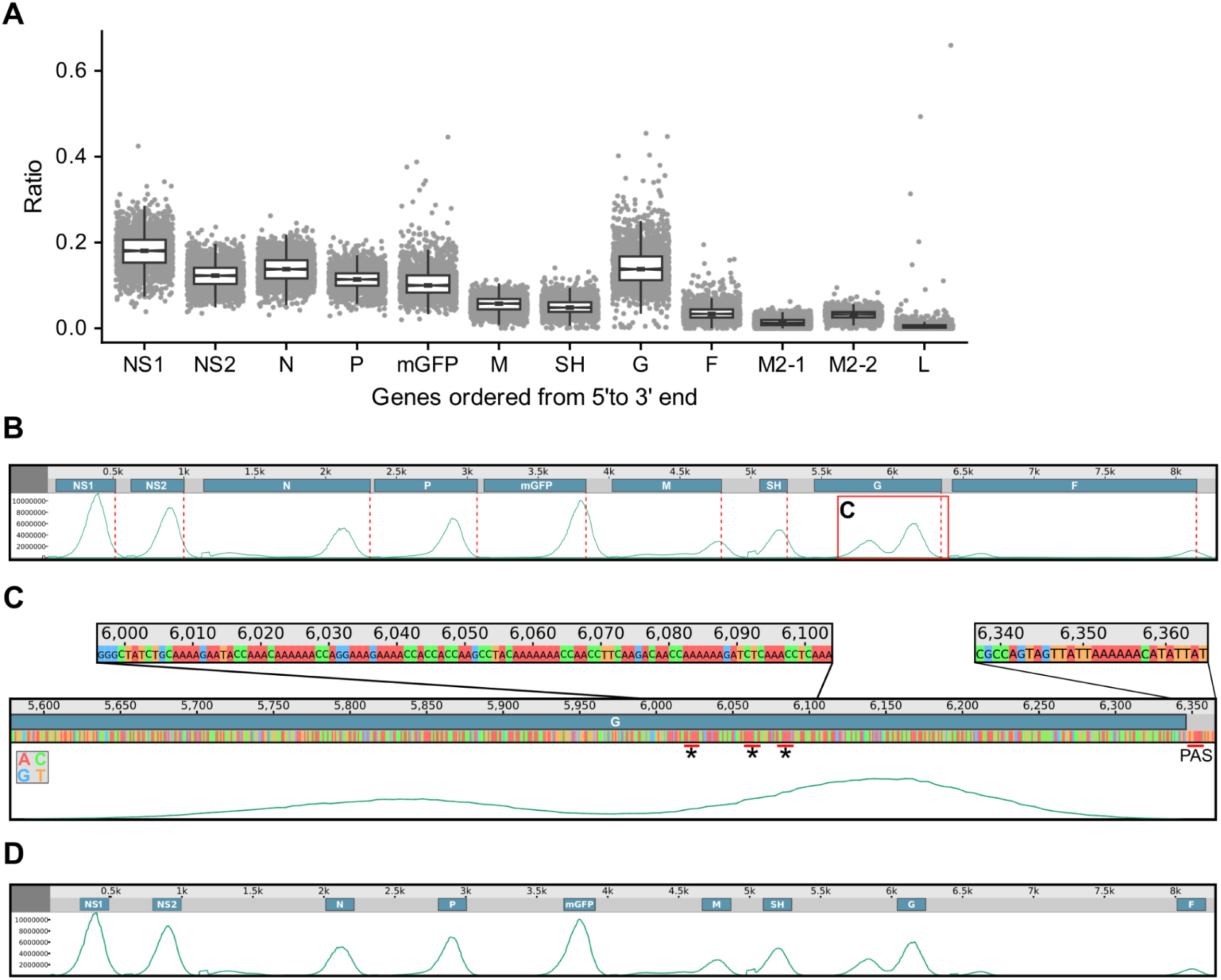
Analysis of reads mapped to RSV genome and reannotation of RSV genome. **(A)** Dot plot of fraction of reads mapped to viral mRNAs with superimposed box plot (XdescriptionX) before manual RSV reannotation. **(B)** Genome Viewer of the RSV genome annotation. Dashed red lines mark the end of annotated genes. Red box labeled ‘C’ of G segment indicates the part of the genome analysed in panel C in detail. **(C)** Genome viewer magnified on the G segment with the polyA signal and several polyA-like signal sequences are highlighted by an asterisk and magnified. **(D)** Genome viewer of the manual RSV genome annotation covering 200 bp windows centered on the 3’ read count peaks.

**Supplementary Figure S5:**
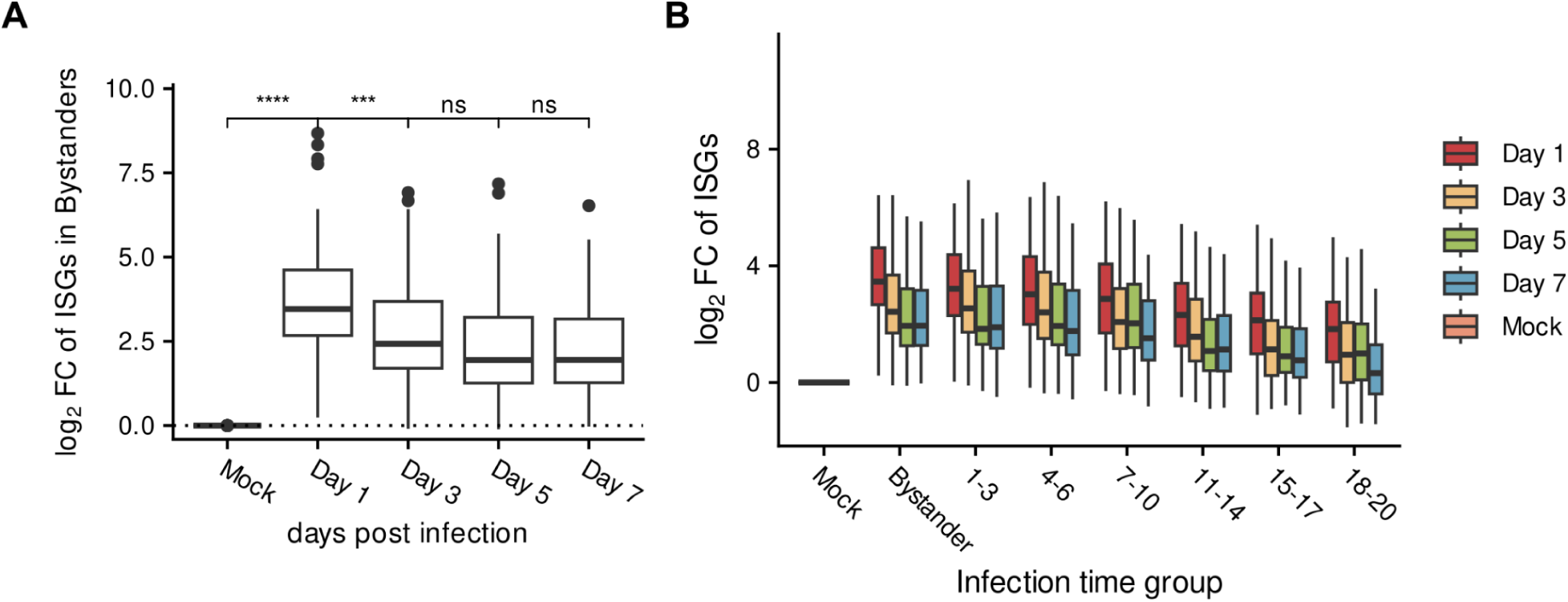
Expression of ISGs across time points. (A) Boxplots showing the log2 fold changes of ISGs across all time points versus all mock cells. (B) Boxplots showing the log2 fold changes of bystander cells and infection time groups and separated by time points.

**Supplementary Figure S6:**
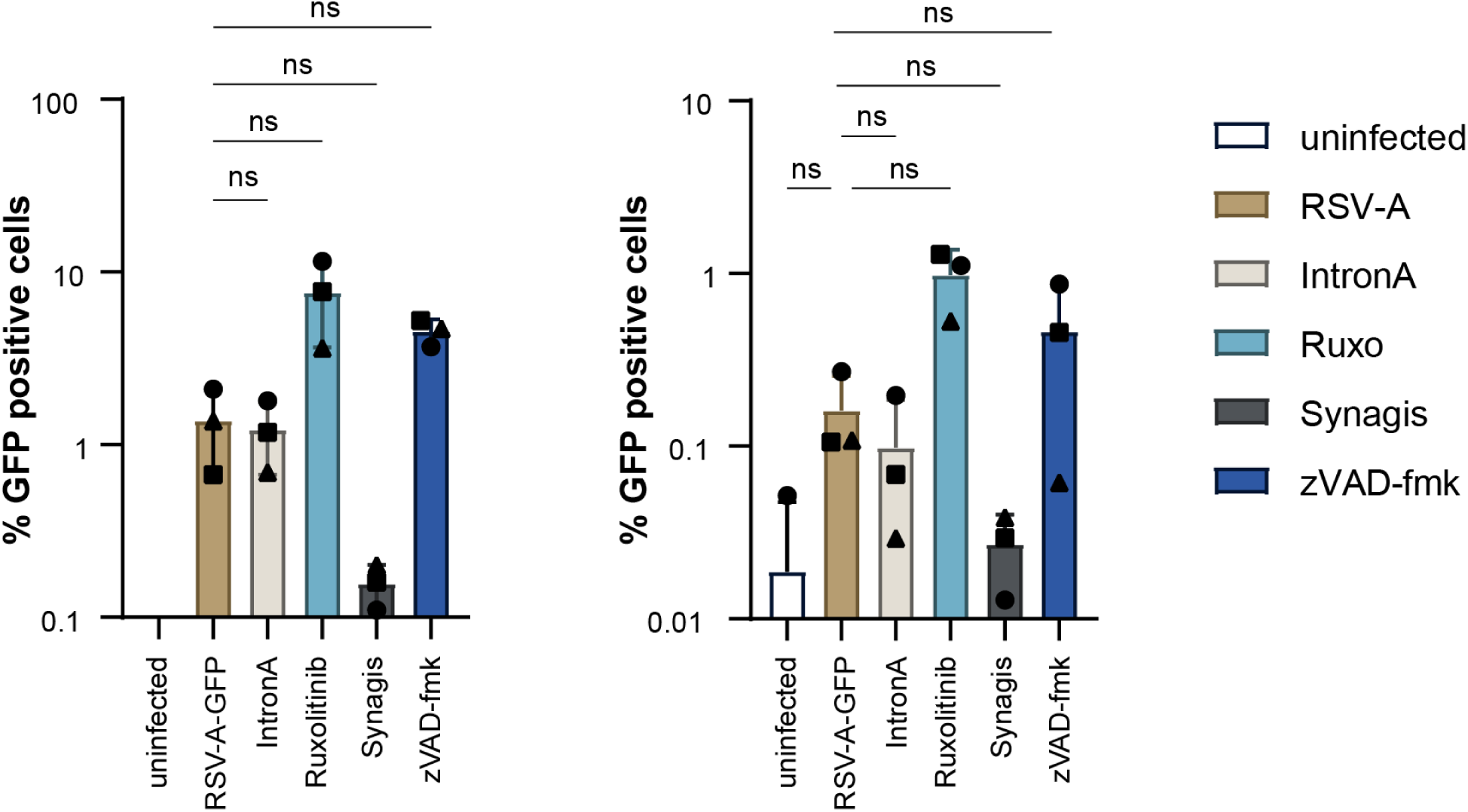
Effect of drug-treatment on RSV infection in primary cells. Well-differentiated primary human airway epithelial cells were inoculated for one hour with an RSV-A-GFP reporter virus. 24h post inoculation, cells were treated with 1000 IU/mL IntronA, 10 µM Ruxolitinib, 10 µg/mL Synagis or 10 µM zVAD-fmk from both sides. To maintain air-liquid interface cultures, the drugs from the apical compartment were removed after 1h and apical treatment was repeated daily while basolateral treatment continued throughout the experiment. 120h post infection, cells were trypsinized and analyzed by flow cytometry using a SONY spectral analyzer SA3800. Experiments were performed twice, each with 3 independent lung cell donors.

**Supplementary Figure S7:**
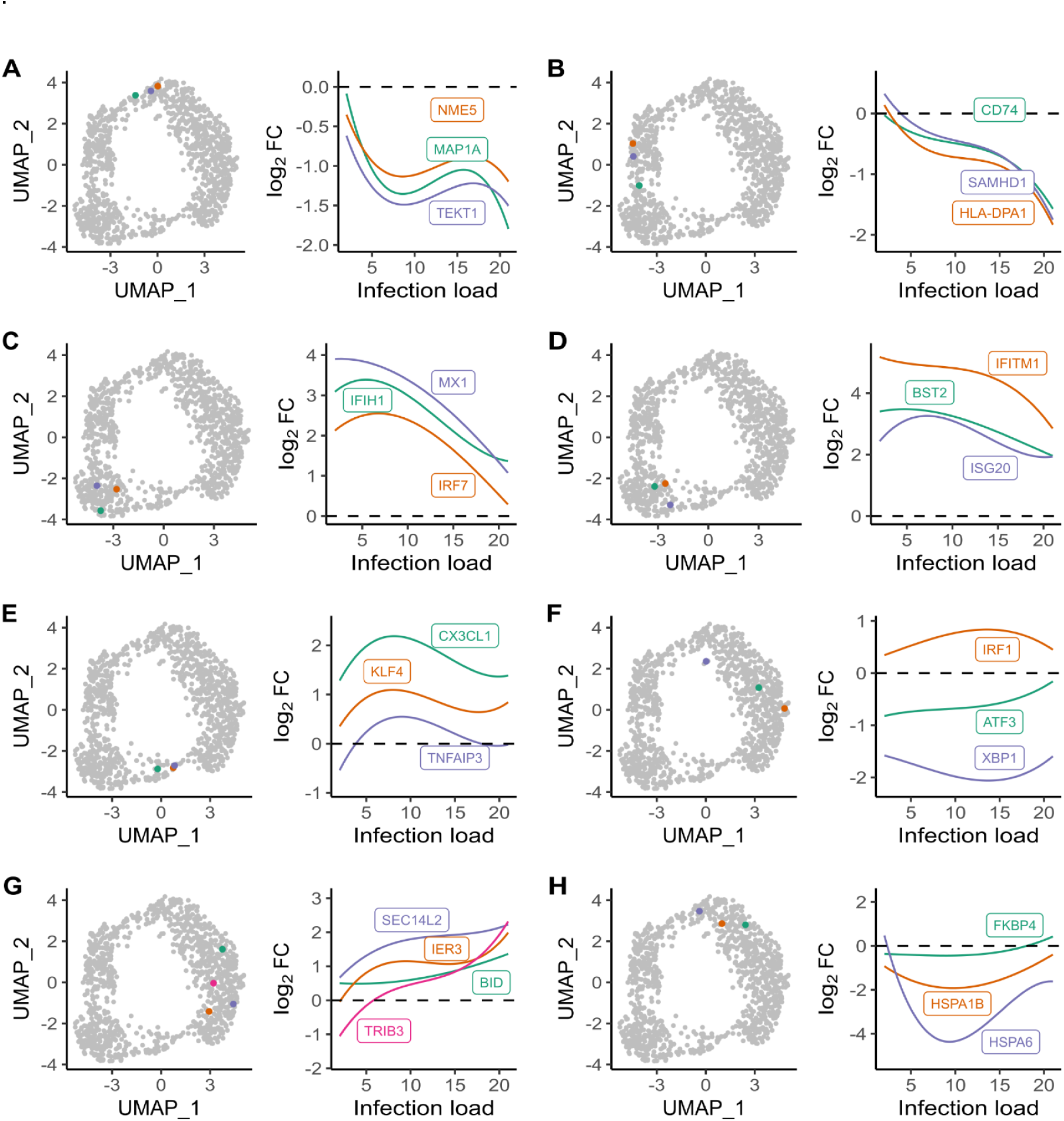
Visualization of Exemplary Genes in Gene Clustering Analysis. **(A-H, left)** UMAP representation of all differentially expressed genes over the Infection time, based on their log2FC values in all pseudobulks. Exemplary genes for every cluster are highlighted. **(A-H, right)** Line plots of mean log2FC values versus Mock over infection time for exemplary genes per cluster.

**Supplementary Figure S8:**
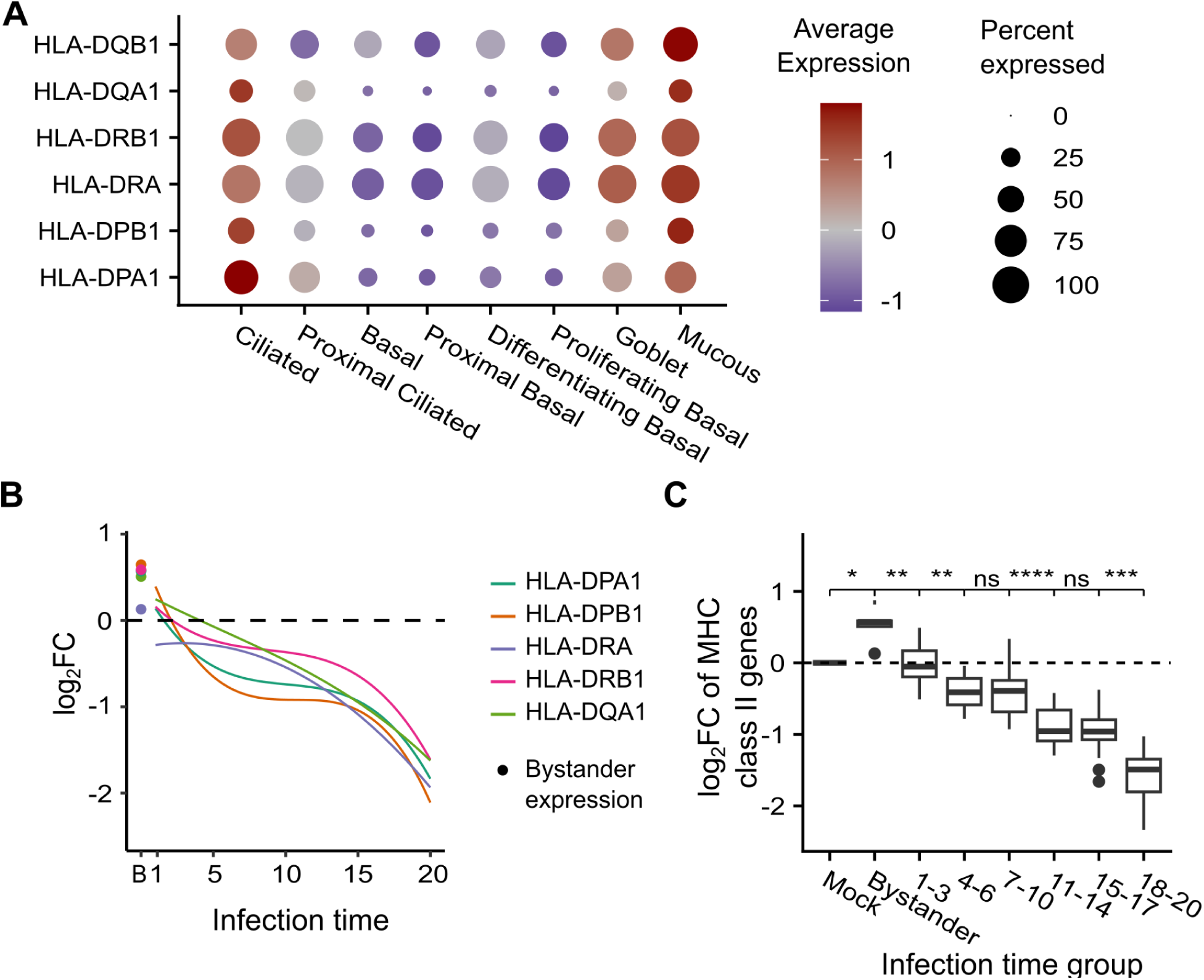
Analysis of HLA genes expression. **(A)** Average expression and percentage of expressing population of MHC class II genes in mock over all cell types. **(B)** Mean LogFC values of differentially expressed genes versus Mock in bystander and infected cells over infection time. HLA-DQB1 was not differentially expressed. **(C)** Boxplots of the mean LogFC of differentially expressed MHC class II genes in mock, bystander and infected cells over Infection load. Wilcoxon test, p-values indicated.

**Supplementary Figure S9:**
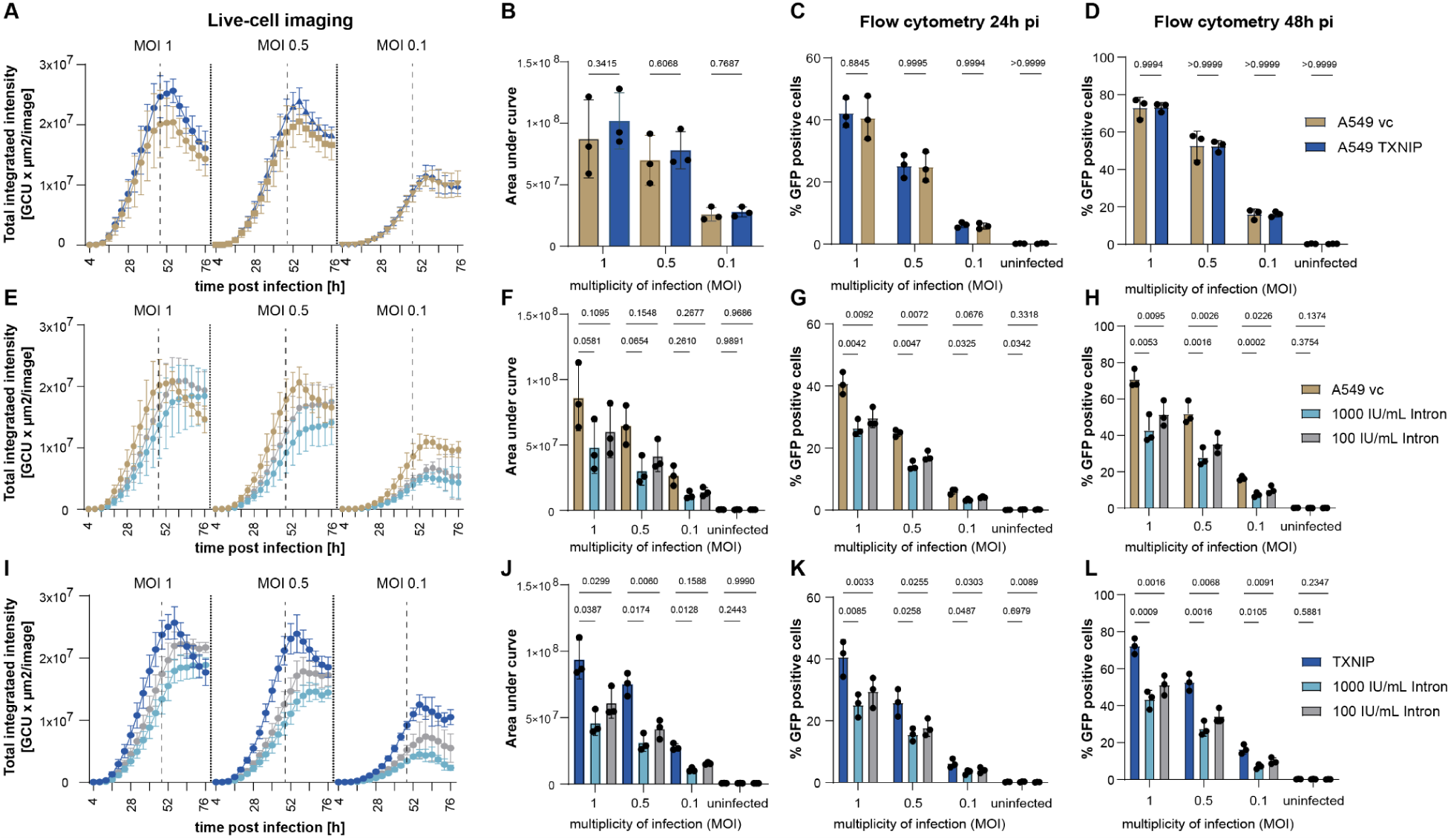
Influence of TXNIP overexpression on RSV infection. (A) A549 vector control (brown) or A549 cells stably overexpressing TXNIP (blue) were inoculated with an RSV-A-GFP reporter virus at indicated multiplicities of infection (MOI) for 3h. Viral inoculum was removed and live-cell imaging using an Incucyte started 4h post infection (pi) at a 10x magnification. Four pictures per well were taken every 4h for up to 76h pi. Total integrated green object intensity (GCU× µm^2^/image) was analyzed using the cell-by-cell software of Sartorius. (B) Area under the curve from (A) until 48h pi (dashed line) was calculated and the statistical analysis was performed using a repeated-measurement one-way ANOVA in combination with Sídák’s multiple comparison test. Mean and std dev of n=3 independent experiments (A+B) as well as the results from the single experiments (symbols, B) is given. (C, D) Flow cytometric analysis of RSV-GFP positive cells at 24h (C) and 48h (D) post infection. Bars represent mean and std dev. of n=3 independent experiments. Two-way ANOVA with Sídák’s multiple comparison test. (E-H) A549 vector control cells were pretreated for 16h prior to infection with an RSV-A-GFP reporter virus at indicated MOÍs with Interferon-2 alpha at two different concentrations. (E, F) Live-cell imaging and statistical analysis of n=3-4 independent experiments is given. (G, H) Flow cytometric analysis of RSV-GFP positive cells 24h pi (G) and 48 pi (H) of n=3 independent experiments is depicted. Bars represent mean and standard deviation including the results for each independent experiment (symbol). Repeated-measurement one-way ANOVA with Sídák’s multiple comparison test. (I-L) A549 TXNIP overexpressing cells were treated with 2 different concentrations of interferon-2alpha 16h prior to infection with an RSV-A-GFP reporter virus at different concentrations and viral replication was monitored using live-cell imaging (I). (J) Area under the curve calculations until 48h post inoculation and statistical analysis of n=3 independent experiments using repeated-measurement one-way ANOVA with Sídáks multiple comparison test. (M) TXNIP protein expression of vector control cells or cells stably transduced with TXNIP.

